# Generation of a versatile BiFC ORFeome library for analyzing protein-protein interactions in live *Drosophila*

**DOI:** 10.1101/343483

**Authors:** J. Bischof, M. Duffraisse, E. Furger, L. Ajuria, G. Giraud, S. Vanderperre, R. Paul, M. Björklund, D. Ahr, A.W. Ahmed, L. Spinelli, C. Brun, K. Basler, S. Merabet

**Author notes:** These authors contributed equally to this work (J. Bischof, M. Duffraisse, E. Furger, L. Ajuria).

## Abstract

Transcription factors achieve specificity by establishing intricate interaction networks that will change depending on the cell context. Capturing these interactions in live condition is however a challenging issue that requires sensitive and non-invasive methods. We present a set of fly lines, called “multicolor BiFC library”, which covers most of the *Drosophila* transcription factors for performing Bimolecular Fluorescence Complementation (BiFC). The multicolor BiFC library can be used to probe binary or tripartite interactions and is compatible for large-scale interaction screens. The library can also be coupled with established *Drosophila* genetic resources to analyze interactions in the developmentally relevant expression domain of each protein partner. We provide proof of principle experiments of these various applications, using Hox proteins in the live *Drosophila* embryo as a case study. Overall this novel collection of ready-to-use fly lines constitutes an unprecedented genetic toolbox for the identification and analysis of protein-protein interactions *in vivo*.

## Introduction

Proteins are distributed in various compartments within the cell, acting in a crowded environment and establishing hundreds of molecular contacts that will eventually dictate cellular function. These molecular contacts are often highly dynamic, may be of weak affinity, and depend on the cell context. Capturing these versatile interactions is therefore a key challenge to better understand the molecular cues underlying protein function *in vivo*.

Two types of experimental strategies are classically used for high-throughput screening of protein-protein interactions (PPIs): yeast two-hybrid (Y2H) and tandem affinity purification coupled to mass spectrometry (TAP-MS). Y2H detects PPIs in an automatable way in live yeast cells, while TAP-MS is based on co-immunoprecipitation (co-IP) and subsequent MS analysis of the different constituents of the complex (see ^1,2^ for review). Recent developments such as proximity based biotinylation (BioID) have significantly increased the sensitivity of the TAP-MS approach, allowing capturing low affinity PPIs from a small number of cells ^3^. Despite their wide range of applications, Y2H and TAP-MS still have several drawbacks. For example, Y2H does not reproduce the plant or animal cell environment, and each revealed interaction needs thus to be validated in the relevant physiological context afterwards. TAP-MS can be performed in the appropriate cell type, but the extraction protocol requires experimental conditions (especially for cell fixation and/or cell lysis) that are often not neutral to the integrity of endogenous PPIs. Finally, BioID allows interaction analyses without fixation, but may capture proteins promiscuously based on proximity rather than direct physical interactions, which could yield to a number of false positives.

In addition to these high-throughput approaches, PPIs can also be analyzed at a low-scale level, for example to validate the interaction status of one protein with few candidate partners. Co-IP followed by western blot is classically used for this purpose. However, this approach requires tools that are not systematically available, such as good antibodies or tagged-constructs. Co-IP experiments may also be poorly sensitive, detecting mainly interactions that will resist cell lysis and purification conditions.

An alternative and more sensitive approach is *in situ* proximity ligation assay (PLA), which provides a direct readout of the candidate PPI in the cell ^4^. A major limitation of PLA is the need of good antibodies against both interacting proteins. In addition, PLA works on fixed material, which is not neutral for PPIs.

Among the few methods that are compatible for PPI analysis in live conditions is Förster (or Fluorescence) resonance energy transfer (FRET), which relies on the transfer of a virtual photon between two fluorescent chromophores upon excitation. This transfer occurs within a small distance (less than 10nm) and can thus be used to validate the close proximity between two candidate proteins, or to capture a conformational change ^5^. FRET requires high level of protein expression and dedicated interfaces to interpret the few emitted signals with confidence and therefore cannot be used for large-scale applications.

By contrast, Bimolecular Fluorescence Complementation (BiFC) appears much more convenient since it is based on a visible fluorescent signal. BiFC relies on the property of monomeric fluorescent proteins to be reconstituted from two sub-fragments upon spatial proximity (in a similar range of distance as FRET). This method has been used in different plant and animal model systems and with various types of proteins ^6–8^. In particular, recent work has established experimental parameters for performing BiFC in the live *Drosophila* embryo ^9^, and the method was coupled to a candidate gene approach to identify new interacting partners of *Drosophila* Hox proteins ^10^.

Moreover, the bright intrinsic fluorescence of BiFC allows analysis of PPIs using commonly available fluorescent microscopes and with normal protein expression levels. Originally established with the Green Fluorescent Protein (GFP ^11^), BiFC has by now been developed with various GFP-derivatives such as the YFP, Venus or Cerulean proteins ^12–14^. BiFC has also been established with other types of monomeric fluorescent proteins, including red fluorescent variants like mRFP1 ^15^ or mCherry ^16^, and more recently the near infrared fluorescent protein iRFP ^17^. In all cases, the complementation between the two sub-fragments of the fluorescent protein induces the formation of covalent junctions, leading to a stabilization of the protein complex. While this property forbids monitoring temporal dynamics of PPIs, this practically irreversible nature of the complementation allows detection of weak and otherwise transient PPIs, making BiFC a very sensitive approach for studying PPIs *in vivo*. BiFC has also been used in several high throughput approaches in yeast ^18^, plant ^19^ and mammalian cells ^20^, or for drug discovery against a specific PPI ^21,22^, demonstrating its suitability for large-scale applications.

Here we present a genetic repertoire covering 453 *Drosophila* transcription factors (TFs) (corresponding to around 65% of annotated TFs) for performing BiFC in a tissue- and developmental stage-specific manner *in vivo*. This genetic repertoire is called multicolor BiFC library and consists of a collection of fly lines that complement the previously established FlyORF collection (https://flyorf.ch/ and ^23^). The multicolor BiFC library is continually updated and aims at covering all *Drosophila* TFs in the near future. We provide proof of concept experiments showing the suitability of the BiFC-FlyORF library for performing either large-scale interaction screens or for analyzing individual PPIs. This collection of fly lines constitutes the first genetic toolbox for analyzing thousands of different PPIs in a live animal organism, opening new avenues for understanding molecular properties of protein interaction networks.

## Results

### Generation of a fly library containing Gal4-inducible ORFs compatible with Venus-based BiFC

560 open-reading frames (ORFs) among the 3000 actually present in the FlyORF library code for TFs. These ORFs are under the control of upstream activation sequences (UAS sites) and fused in frame to a hemagglutinin tag (3xHA) sequence at their 3’ end ^23^. The ORFs are flanked by distinct FRT sites that can be used to replace the promoter sequence and/or the 3xHA tag region by any other sequence of choice upon FLP/FRT-mediated recombination *in vivo* ^23^. In particular, two swapping fly lines have been generated to replace the C-terminal 3xHA-tag by sequences coding for the N- or C-terminal fragment of the Venus fluorescent protein (fragments respectively called VN and VC hereafter in the manuscript ^23^). These two fragments, when attached to proteins that are co-expressed, are able to complement upon spatial proximity, allowing assessment of the interaction by BiFC (Figure 1A). A proof of principle experiment between two known interacting partners present in the FlyORF library proved the efficiency of the swapping and the specificity of the Venus-based BiFC in the *Drosophila* wing imaginal disc ^23^.

**Figure 1.**
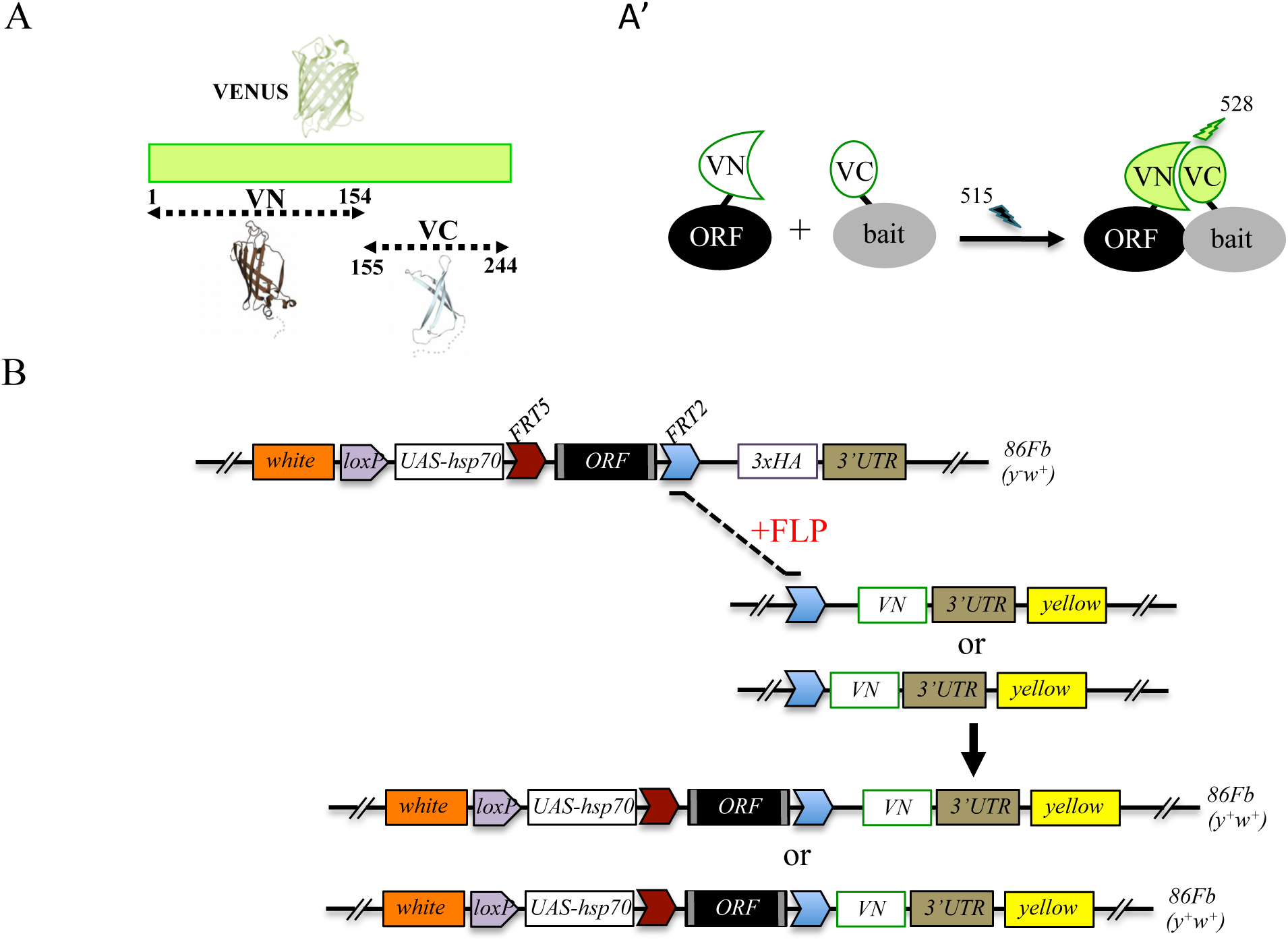
Generation of a Gal4 inducible library compatible with Venus-based BiFC in *Drosophila.* **A-A’.** Principle of the Venus-based BiFC between a candidate ORF (Open Reading Frame) and a bait protein fused to the N- (VN) or C-terminal (VC) fragment of Venus, respectively. Excitation and emission wavelengths are indicated. **B.** Principles of Flippase (FLP)/FRT-mediated recombination to swap the C-terminal 3xHA tag of the ORF with the original VN or new VN-short tag line. Genetic crosses and selection procedure are described in ^23^. Note that the UAS-ORF-HA and resulting UAS-ORF-VN are located on the third chromosome (*86Fb*). See also Supplementary Figure 1 and Table S1.

Experimental parameters for performing Venus-based BiFC have also been established in different tissues of the live *Drosophila* embryo ^9^. In particular, several controls showed that Venus-based BiFC could not occur in conditions where the interaction between the two candidate partners was strongly affected. This was demonstrated by using mutant proteins ^9^ or by coexpressing one partner that could not complement (not fused to a Venus fragment) and thus competes against BiFC ^10^.

Although very convenient for generating new fusions without additional fly transgenesis, swapping experiments require several generations, and therefore several weeks, before getting the desired BiFC-compatible fly line ^24^. In order to introduce ready-to-use fly lines for performing Venus-based BiFC, we systematically exchanged the 3xHA tag in a number of TF lines of the FlyORF library for the VN tag (Figure 1B). These first rounds of swapping experiments led to a collection of 137 ORF-VN fly lines (Table S1, third column). We further generated an alternative VN-swapping fly line that allows fusing the ORF to the Venus fragment, however with a shorter linker region between ORF and VN (Figure 1B and Supplementary Figure 1, see also Materials and Methods). Reducing the length of this region could potentially diminish the risk of revealing indirect PPIs *in vivo*. A series of pilot tests confirmed that the new VN-short tag fly line is suitable for swapping and BiFC experiments (Supplementary Figure 1). This new swapping fly line is now systematically used for generating additional ready-to-use ORF-VN tagged fly lines. To date, 73 ORFs have been fused to the VN fragment with the short linker region (Table S1, fourth column). In addition to the previously generated fly lines (including those generated in ^9,25^, Table S1, fifth column), the multicolor BiFC library currently contains 235 different TFs fused to VN for doing Venus-based BiFC in *Drosophila*.

### Generation of a fly library containing Gal4-inducible ORFs compatible with bicolor BiFC

One interesting feature of BiFC is the opportunity of using fragments from various GFP-derived proteins to visualize two different PPIs in the same cell ^13^. In particular, it was shown that the C-terminal fragment of the blue fluorescent Cerulean protein (CC) could complement with either the N-terminal fragment of Venus (VN) or Cerulean (CN), giving rise to Venus or Cerulean-like fluorescent signals, respectively (Figure 2A-A’ and ^13^). Given that Cerulean-based BiFC was shown to be exploitable in the *Drosophila* embryo ^9^, we decided to increase the versatility of the BiFC library by generating an additional set of fly lines that could be used for both Venus- and Cerulean-based BiFC (Figure 2B). This new UAS-inducible ORF-CC library was inserted on the second chromosome (see Materials and Methods and Supplementary Figure 2) and can be used to perform Venus- and Cerulean-based BiFC with any VN- (including VN fusion constructs of the library) and CN-fused bait protein, respectively. Fly lines for 326 different ORF-CC have been generated so far (Table S2).

**Figure 2.**
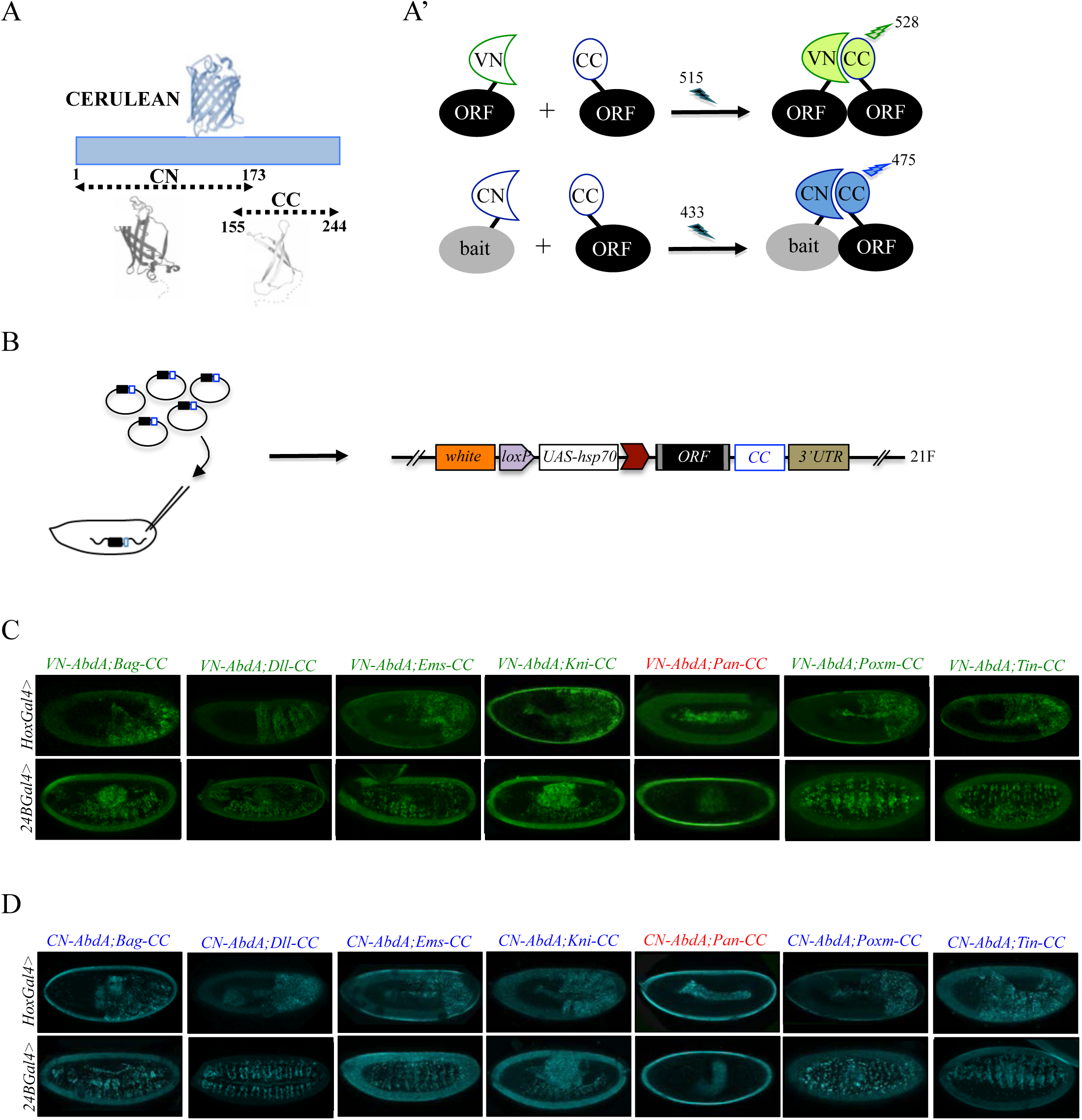
Generation of a Gal4 inducible library compatible with Venus- and Cerulean-based BiFC in *Drosophila*. **A-A’.** Principle of bicolour BiFC by using the complementation property between the C-terminal fragment of the blue fluorescent protein Cerulean (CC) and the N-terminal fragment of Venus (VN) or Cerulean (CN). Excitation and emission wavelengths are indicated. **B.** Principle of the generation of the UAS-ORF-CC library at the 21F genomic locus. See also Materials and Methods. **C-D.** Illustrative confocal captures of Venus-based BiFC obtained from different ORF-CCs and VN-AbdA (C) or CN-AbdA (D) interaction partners, as indicated. Fusion proteins are expressed with the *abdA-Gal4* (upper panels) or *24B-Gal4* (lower panels) driver and BiFC is observed in the epidermis (stage 10/11) or somatic mesoderm (stage 14), respectively. Note that the amnioserosa, the gut inside the embryo and the vitelline membrane around the embryo display strong autofluorescence. Absence of interaction with Pangolin (Pan) is highlighted in red. See also Supplementary Figures 2-4 and Tables S2 and S3.

To verify that the ORF-CC library is compatible with Cerulean- or Venus-based BiFC, we chose a set of six TFs that have already been analyzed as VN-fusion constructs with the Hox protein AbdominalA (AbdA) fused to the VC complementary fragment (Bagpipe, Bap; Empty spiracles, Ems; Knirps, Kni; Pangolin, Pan; Pox mesoderm, Poxm; Tinman, Tin), and whose interaction status was also validated by co-IP experiments ^10^. Among those six TFs, all but Pan could interact with AbdA ^10^. The same pool of TFs was here tested as ORF-CC fusions with VN-AbdA or CN-AbdA in two different tissues (epidermis and somatic mesoderm) and at two different developmental stages (stages 10 and 12) by using two different Gal4 drivers. Results showed that Venus and Cerulean fluorescent signals could easily be distinguished from the fluorescent background in the epidermis or mesoderm in the case of all tested TFs except Pan (Figure 2C-D). Moreover, competition tests with the corresponding HA-tagged ORFs validated the specificity of BiFC obtained between VN-AbdA and each candidate ORF-CC construct (Supplementary Figure 3). This last experiment confirms that the ORF-3xHA fly lines from the original FlyORF library can be used to verify the specificity of BiFC signals. Altogether, these observations establish that the ORF-CC constructs of the multicolor BiFC library are compatible with either Cerulean- or Venus-based BiFC in different tissues of the live *Drosophila* embryo.

### Using the multicolor BiFC library for large-scale interaction screens in live *Drosophila* embryos

The multicolor BiFC library currently covers 453 different *Drosophila* TFs (around 65% of annotated TFs), making a total of 579 fly lines due to several TFs fused with different VN and/or CC versions. Among the 453 TFs, 127 are available as VN fusion constructs, 219 as CC fusion constructs, and 108 as VN- and CC-fusion constructs (Supplementary Figure 4). Together with the simplicity of genetic crosses and readouts, it makes this library appropriate for large-scale interaction screens in *Drosophila*.

We provide a proof of concept by analyzing interaction properties of 260 TFs of the multicolor BiFC library with Ultrabithorax (Ubx) and AbdA. These two Hox proteins have highly similar domains and motifs (Figure 3A). They share a number of common functions during embryogenesis and can substitute for each other in several tissues, as noticed for example in the epidermis ^26^, trachea ^27^ or ventral nerve cord ^28^. Ubx and AbdA have also few distinct expression patterns that correlate with specific functions in the embryo. For example, the specification of oenocytes ^29^, heart ^30^ or gonadal mesoderm ^31^ is under the control of AbdA, a function that cannot be substituted by Ubx. Such exclusive functions are however rare during *Drosophila* embryogenesis, suggesting that most embryonic functions of Ubx and AbdA could rely on the interaction with a large number of common cofactors. In this context, we aimed at using the multicolor BiFC library to reveal interactions that could be specific of AbdA. The identification of such partners would validate the specificity and sensitivity of the multicolor BiFC library in the context of a large-scale interaction screen with two closely related bait proteins.

**Figure 3.**
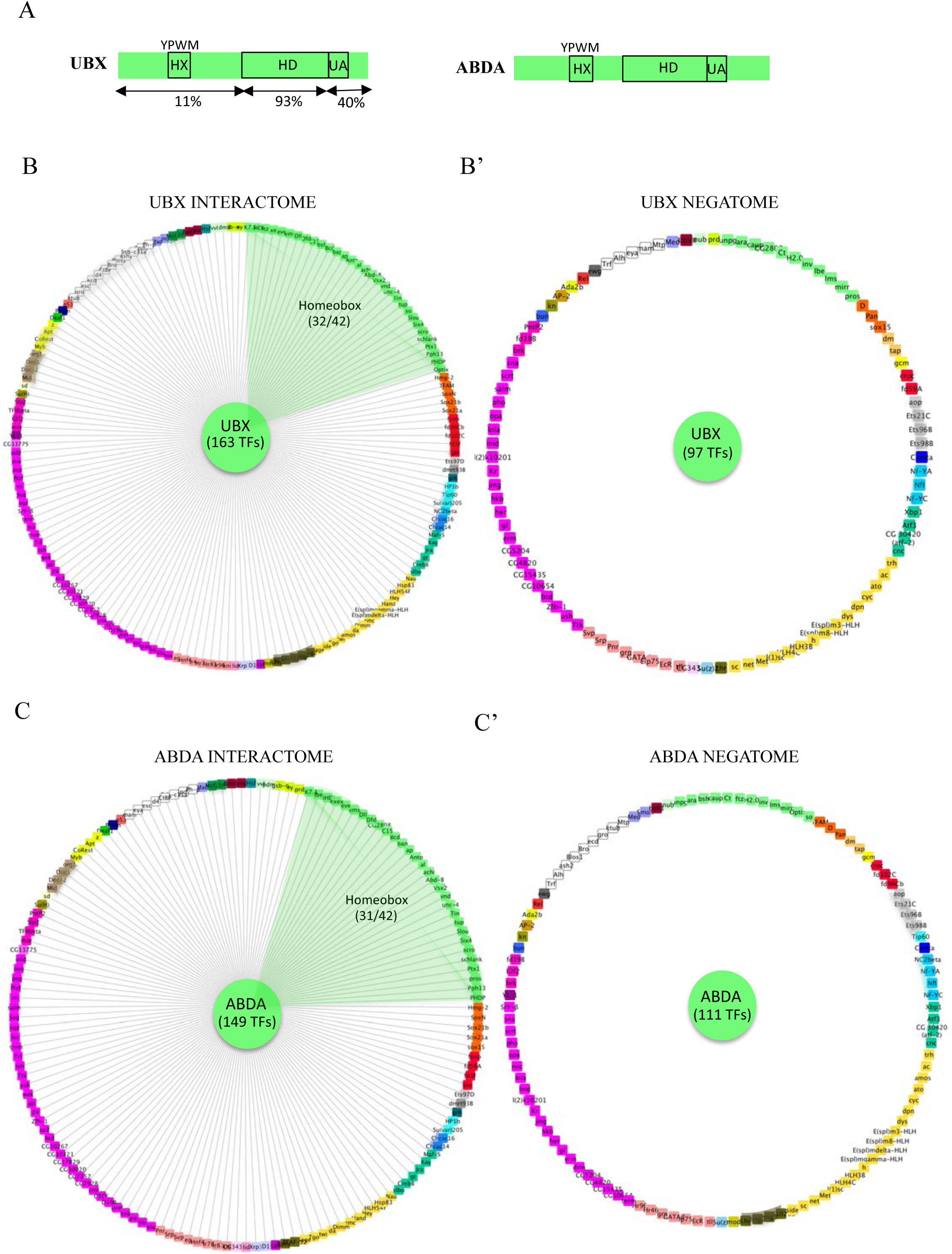
Using the multicolor BiFC library to reveal interactomes and negatomes of Ubx and AbdA in the live *Drosophila* embryo. **A.** Schematic representation of Ubx and AbdA. The homeodomain (HD), hexapeptide (HX) and UbdA (UA) motifs are indicated, as well as the percentage of identity of Ubx with AbdA along the corresponding regions. **B-C.** Representation of the interactome of Ubx (B) or AbdA (C). **B’-C’.** Representation of the negatome of Ubx (B’) or AbdA (C’). Each color code corresponds to a different family of transcription factors (TFs). Only homeobox-containing TFs are significantly enriched in the Ubx and AbdA interactomes (highlighted in green). Networks are represented using Cytoscape 3.6 ^56^. See also Supplementary Figures 5-12 and Tables S4-S6 and S9.

Among the 260 different TFs, 127 were analyzed in fusion with VN and 133 in fusion with CC. Thirty-five TFs were tested in fusion with VN and CC to assess the influence of the fusion topology on BiFC results (Table S9, see also Discussion). Interactions were analyzed in the epidermis of stage 10 embryos, in the *Ubx*- or *abdA*-expression domain, by using *Ubx-Gal4* or *abdA-Gal4* driver, as previously described ^10^.

The analysis showed that 62% (163/260) and 57% (149/260) of the tested TFs could interact with Ubx (Figure 3B, Supplementary Figures 5-6 and Table S4) or AbdA (Figure 3C, Supplementary Figures 7-9 and Table S4), respectively. Among all positive interactions, 50% (130/260) are common to the Ubx and AbdA interactomes (Supplementary Figure 10), consistent with the numerous overlapping functions between the two Hox proteins during embryogenesis. Interestingly, the homeodomain (HD) class of TFs appears significantly enriched in the Ubx (p value=0,041) and AbdA (p value=0,017) interactomes, suggesting that HD-containing TFs could constitute a privileged class of cofactors (Figure 3B-C). We also looked at TFs that did not interact with the Hox proteins, constituting the so-called negatomes ^32^, but did not notice enrichment for any particular class of TFs (Figure 3B’-C’). Overall, the BiFC screen shows that the two Hox proteins could interact with a surprisingly high number of various types of TFs. A similar conclusion (with 41% of positive interactions) was obtained from a previous candidate gene screen based on competitive BiFC with a set of 80 TFs ^10^. This high interaction potential of Hox proteins could be explained by their numerous functions during embryogenesis and the extreme sensitivity of BiFC (linked to the fluorescent signal and UAS/Gal4 expression system). Such a high rate of positive interactions necessitates further development of alternative approaches to validate the specificity of our BiFC observations. In the following, we describe additional experimental strategies that revealed new features of Ubx and AbdA interactomes and confirmed that interactions observed with BiFC are of functional significance.

The BiFC screen was performed in the *Ubx-* or *abdA*-expression domain by expressing the TFs using *Ubx-Gal4* or *abdA-Gal4* driver. True positive interactions are expected to occur between TFs that are co-expressed in the same spatial expression domain. To test this, we considered the extent of the overlap between the expression domain of Ubx or AbdA and each TF (as the ratio between the number of tissues in which the TF and Ubx or AbdA are co-expressed and the total number of tissues composing the TF expression domain during embryogenesis). To this end, we used annotations from ^33^ and the Flybase database to assign the expression status of Ubx, AbdA and each TF in 25 different developmental contexts (Table S4). This analysis showed that the extent of co-expression is significantly higher when TFs and Hox protein interactions are detected with BiFC than when they are not (Wilcoxon test, pvalue=1.10^−2^ and 5.10^−5^ for Ubx and AbdA, respectively: Supplementary Figure 11). Thus, the BiFC screen reveals more frequently interactions between TFs and Hox proteins when they are co-expressed during embryogenesis.

### The multicolor BiFC library reveals Hox-specific interactomes with distinct enrichments in predicted protein disorder and short linear motifs

Although Ubx and AbdA shared a number of common interactions, our BiFC screen also revealed interactions that were specific for only one of the two Hox proteins. These interactions represent 20% and 13% of their overall interactome, respectively (33 TFs for Ubx and 19 TFs for AbdA, Supplementary Figure 12). This result confirms that the multicolor BiFC library can be used to capture specific interaction partners between two closely related bait proteins.

To get more insights into the molecular properties of Ubx- and AbdA-specific interactomes, we analyzed their overall content in intrinsically disordered regions (IDRs) and short linear motifs (SLiMs, see Material and Methods), which are key molecular features underlying PPIs *in vivo* ^34^. Interestingly, we observed that AbdA-specific interactors are strongly enriched in predicted IDRs and SLiMs, while Ubx-specific interactors have few SLiMs and are enriched in ordered domains (Table S5). This molecular distinction does not appear in the common Hox interactome or negatome, further arguing that the screen identified true positive interactions (Table S5). Thus, Ubx- and AbdA-specific interactomes contain TFs that display opposite enrichments in their IDRs and SLiMs content.

Further analysis was focused on the AbdA-specific interactome. We noticed that this interactome was enriched in TFs expressed in the fat body/gonad primordium (p value=8.10^−4^) when compared to all positive interactions (encircled TFs in Figure 4B, and Table S6). In contrast, TFs expressed in this tissue are not present in the Ubx-specific interactome. This observation is consistent with the specific role of AbdA in the gonad primordium ^35^. Another TF, Spalt major (Salm), is also found in the AbdA-specific interactome (encircled by a dotted-line in Figure 4B). Among other functions, Salm is important for oenocyte specification ^36^, a role also specifically ensured by AbdA during embryogenesis ^29^. A significant enrichment is also observed in the dorsal vessel (p value=5.10^−2^, TFs annotated with a star in Figure 4B, and Table S6), which again coincides with a specific function of AbdA in particular for the differentiation of ostia and heart beating activity ^30^. By comparison, TFs expressed in tissues where Ubx and AbdA have redundant/common functions show no significant enrichment in the AbdA-specific interactome (i.e epidermis, CNS, somatic mesoderm and trachea: Table S6). Together these observations underline that the multicolor BiFC library is efficient in revealing relevant candidate cofactors involved in AbdA-and tissue-specific functions.

**Figure 4.**
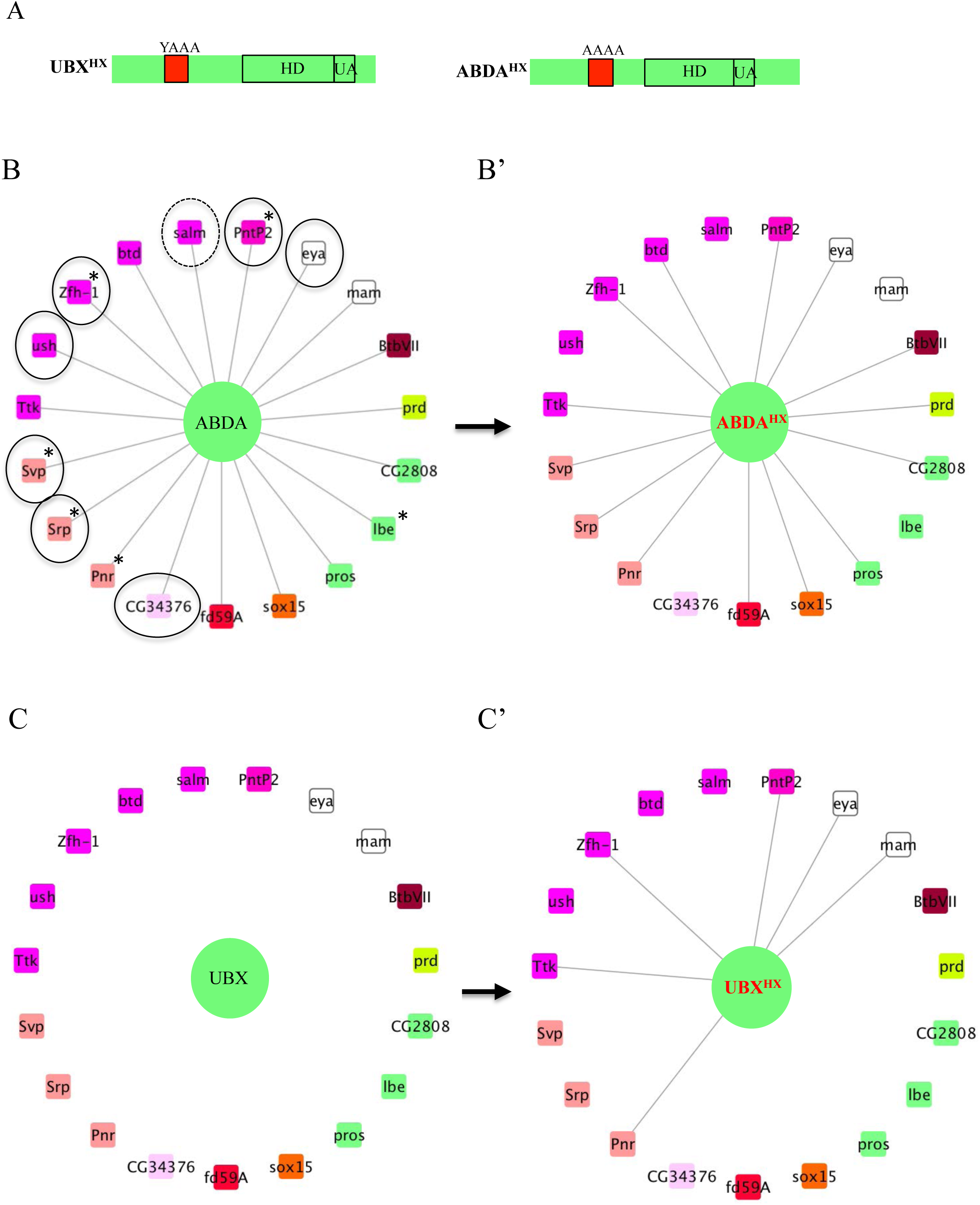
Role of the Hexapeptide (HX) in specifying Ubx and AbdA interactomes. **A.** Schematic representation of HX-mutated Ubx and AbdA proteins. **B.** AbdA-specific interactome. TFs involved in gonad mesoderm or oenocytes specification, which correspond to AbdA-specific functions, are surrounded by a solid or dotted line, respectively. “*” indicates TFs involved in heart specification. **B’**. Effect of the HX mutation in the context of the AbdA-specific interactome. The HX mutation leads to a loss of interaction for 5 TFs (illustrated by the absence of a solid line). **C.** Representation of the AbdA-specific interactome around Ubx. Absence of solid lines illustrates absence of interaction. **C’.** The HX mutation of Ubx induces new interactions with six different TFs (solid black lines).

### A short conserved Hox protein motif controls Ubx- and AbdA-specific interactions

In addition to the presence of characteristic residues within the HD, Hox proteins contain another generic molecular feature that corresponds to a short motif called hexapeptide (HX). This motif is present upstream of the HD in all members of paralog groups 1 to 10 and has primarily been described for its role in mediating the interaction with PBC-class cofactors ^37^. Recent work showed that this motif could also positively or negatively modulate the interaction potential of *Drosophila* Hox proteins with different TFs in the embryo ^25^. Interestingly, this regulatory activity was different depending on the Hox protein and tissue considered ^25^, highlighting that the HX motif is an interface sensitive to the protein environment.

Here we addressed the role of the HX motif in the context of the AbdA-specific interactome (Figure 4A). In particular, the 19 AbdA-specific TFs were analysed with the HX-mutated form of AbdA (Figure 4B). This analysis showed that 5/19 interactions were lost in this context (Figure 4B’). Conversely, the same pool of 19 TFs was also analysed with the HX-mutated form of Ubx (Figure 4A), to assess whether the HX motif of Ubx could also be important for inhibiting specific PPIs. Among this pool of TFs that do not interact with wild type Ubx (Figure 4C), six became positive with HX-mutated Ubx (Figure 4C’). Thus, mutating the HX motif of Ubx relieves an inhibitory activity against specific TFs, reproducing interaction properties of AbdA. In total, dual activities of the HX motif were observed for 11/19 AbdA-specific TFs, highlighting the important role of this motif for specifying Ubx and AbdA interactome properties *in vivo*. More generally, these results show that a common and highly conserved motif between two related Hox proteins acts as a specificity factor, displaying different or even opposite activities for regulating interaction properties with the same pool of TFs.

### Validating BiFC observations by a functional genetics approach in haltere primordium

Since BiFC revealed a number of potential interactions, we asked whether hits from the large-scale interaction screen could somehow be confirmed to contribute to Hox protein function. To this end, we searched for a sensitive genetic background where any subtle modification in Hox protein function could lead to a quantifiable phenotype. In this context, the loss of a Hox cofactor could affect the sensitized Hox protein function and lead to a stronger phenotype. No sensitive Hox-dependent phenotypes are known in the embryo, but several exist in the adult, like eye reduction ^38^ or antenna-to-leg transformation ^39^. These phenotypes rely however on ectopic expression of the Hox product and are therefore not ideal for assessing the role of a candidate cofactor in the normal developmental context. To circumvent this problem, we considered the haltere-to-wing phenotype, which results from the specific loss of Ubx in the haltere primordia ^40^. A particular combination of *Ubx* mutant alleles has more recently been used to measure the ability of different Ubx isoforms to rescue haltere formation ^41^. Importantly, the haltere-to-wing phenotype is sensitive to the dose of Ubx, and removing one copy of the Hox gene is sufficient to induce the formation of few small hairs that are normally found in the wing margin, revealing a weak haltere-to-wing transformation phenotype (Figure 5A-B). In this context, RNAi against *Ubx* is sufficient to induce a strong haltere-to-wing transformation, with the formation of numerous wing-like hairs and a flattened wing-like shape (Figure 5C). This result highlights that the heterozygote *Ubx* mutant phenotype can be increased when *Ubx* function is affected. We decided to use this sensitive background for measuring the role of TFs tested in our BiFC screen (see Materials and Methods). The rational was that affecting the expression of a positive TF acting as a Ubx cofactor in the haltere primordium should increase the haltere-to-wing phenotype. Reversely, a negative TF should have no effect.

**Figure 5.**
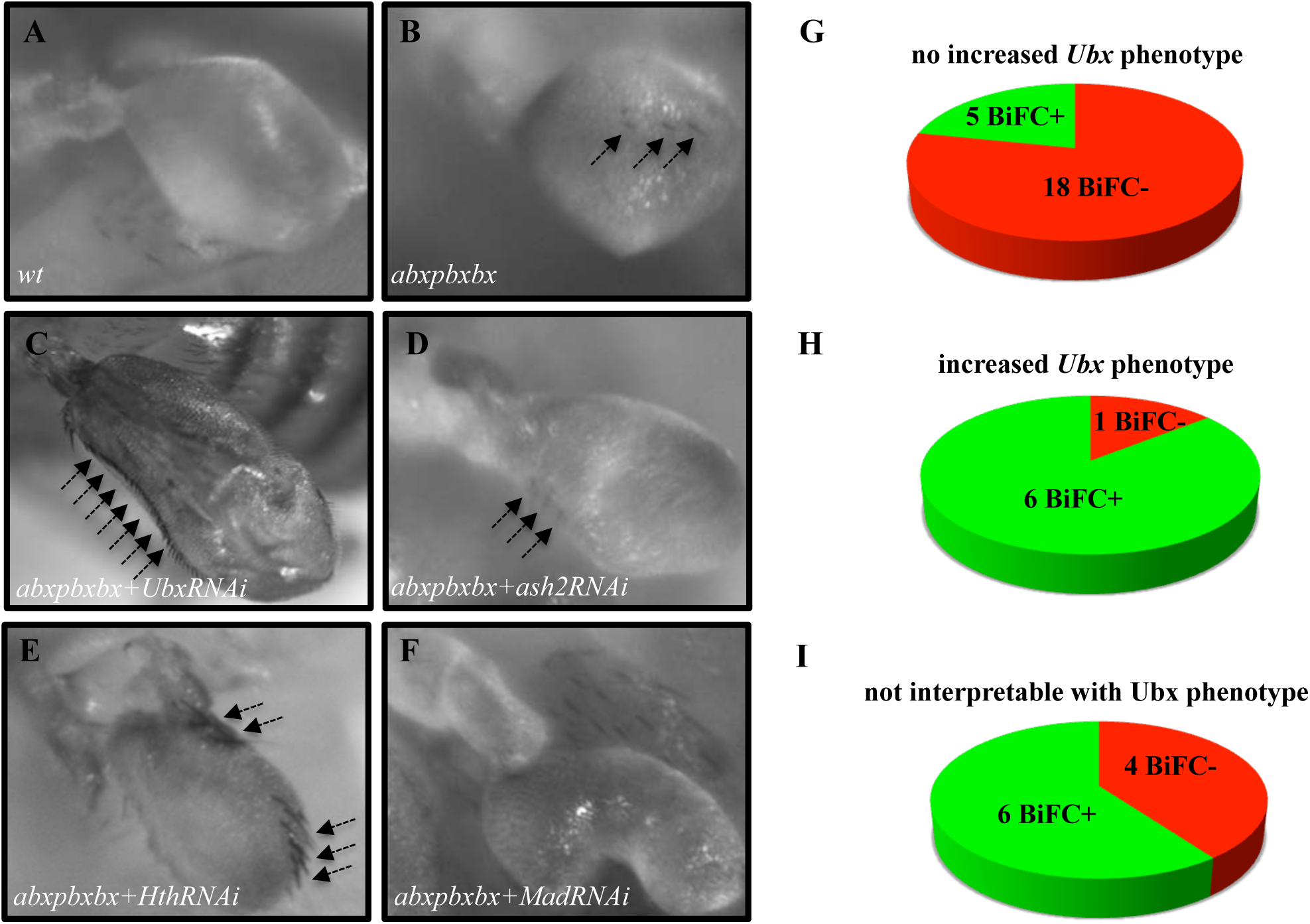
Functional genetics validates BiFC observations with Ubx in haltere primordium. **A-F.** Haltere phenotypes in the different genetic backgrounds, as indicated. Compared to wild type (A), halteres of individuals heterozygous for the Hox regulatory mutation *abxpbxbx* have ectopic short wing-like hairs (arrows in B). This phenotype is increased when affecting *Ubx* expression upon expression of RNAi (arrows in C) or when expressing a RNAi against a TF (shown here for *Homothorax*, *Hth*) that could be required for Ubx function (arrows in E). In contrast, expression of RNAi against a TF that is not required for Ubx function (as shown for *absent_small_or_homeotic_discs _2*, *ash2*) does not increase the phenotype (D). The expression of RNAi against TFs can also affect more globally the haltere (and wing) formation (as shown for Mad), which is difficult to interpret in term of homoeotic transformation and therefore with regard to a potential Ubx cofactor function (F). **G-H.** Diagrams showing the distribution of TFs that were BiFC positive (green) and negative (red) with Ubx in the different cases (not increased haltere phenotype (G); increased haltere phenotype (H); not interpretable (I)). See also Table S7.

TFs were selected based on their known expression and/or function in the haltere (Flybase database, ^42^ and Table S7). The resulting 40 TFs (among all the 260 that were tested in the embryo BiFC screen) were assayed using RNAi experiments in *Ubx* heterozygous mutant haltere discs. Slightly more than half of the tested TFs (23/40: Table S7) did not increase the phenotype of the heterozygous *Ubx* mutant upon RNAi (an illustrative picture is given in Figure 5D). The large majority of those TFs (18/23) were classified as BiFC negative with Ubx in the embryo, confirming that BiFC was specific enough not to identify these as interacting partners of Ubx (Figure 5G and Table S7). In addition, we cannot exclude the possibility that the five other TFs that were BiFC positive could act as Ubx cofactors in another developmental context. Among the 17 remaining TFs, 7 enhanced the phenotype upon RNAi (Table S7). The phenotype enhancement was not as strong as with the RNAi against Ubx, and consisted in the appearance of numerous wing-like hairs in the haltere (an illustrative picture with the strongest phenotype is provided in Figure 5E). 6/7 of those TFs scoring positively in the RNAi assay were also BiFC-positive in the embryo, highlighting that Ubx cofactors were efficiently captured in the BiFC interaction screen (Figure 5H and Table S7). It is also worth mentioning that the TF that gave the strongest phenotype upon RNAi, Homothorax (Hth), displays enriched binding adjacent to Ubx binding sites genome wide in the haltere tissue ^43^. This suggests that Hth could constitute a crucial cofactor for Ubx in the haltere specification program. Finally, RNAi against 10 TFs led to malformations that were difficult to interpret with regard to a potential role as Ubx cofactor since the morphological defects could also be independent of a Ubx cofactor function. An illustrative phenotype is given with a RNAi against the TF Mad (Figure 5I). Because of this ambiguous putative role, these TFs were not further considered (black boxes in Table S7).

Overall, the haltere sensitized genetic background highlighted that 80% (24/30) of the selected TFs that give an interpretable phenotype recapitulated the BiFC observations. The positive or negative BiFC interaction status in the embryo could thus be reproduced at the functional level in another developmental context. This result also suggests that the interaction potential revealed by BiFC with the UAS/Gal4 system is not strongly influenced by the tissue type.

### Using the multicolor BiFC library for analyzing two different PPIs in the same embryo

In addition to large-scale interaction screens, the multicolor BiFC library aims at providing new perspectives for the analysis of individual PPIs *in vivo*. In particular, the ORF-CC library allows performing Venus- and Cerulean-based BiFC, therefore analysing two different PPIs in the same embryo. This so-called “multicolor” property was established in live cells ^13^ and recently extended for use in live *Drosophila* embryos ^9^.

Here we asked whether the multicolor BiFC library could be used for analyzing two different PPIs simultaneously. This potential was more precisely examined in the context of a well-known partnership between AbdA and the cofactor Extradenticle (Exd). Exd belongs to the PBC class of TALE (Three Amino Acids Loop Extention) TFs ^44^. PBC proteins constitute an ancestral and generic class of Hox cofactors, and the activity of Hox-PBC complexes has been described in numerous developmental contexts ^45^. In particular, the partnership between AbdA and Exd is involved in early (e.g., patterning) and late (e.g., gut morphogenesis) developmental processes during *Drosophila* embryogenesis, suggesting that a number of TFs could interact with the two proteins, potentially participating in the activity of AbdA/Exd complexes (Figure 6A).

**Figure 6.**
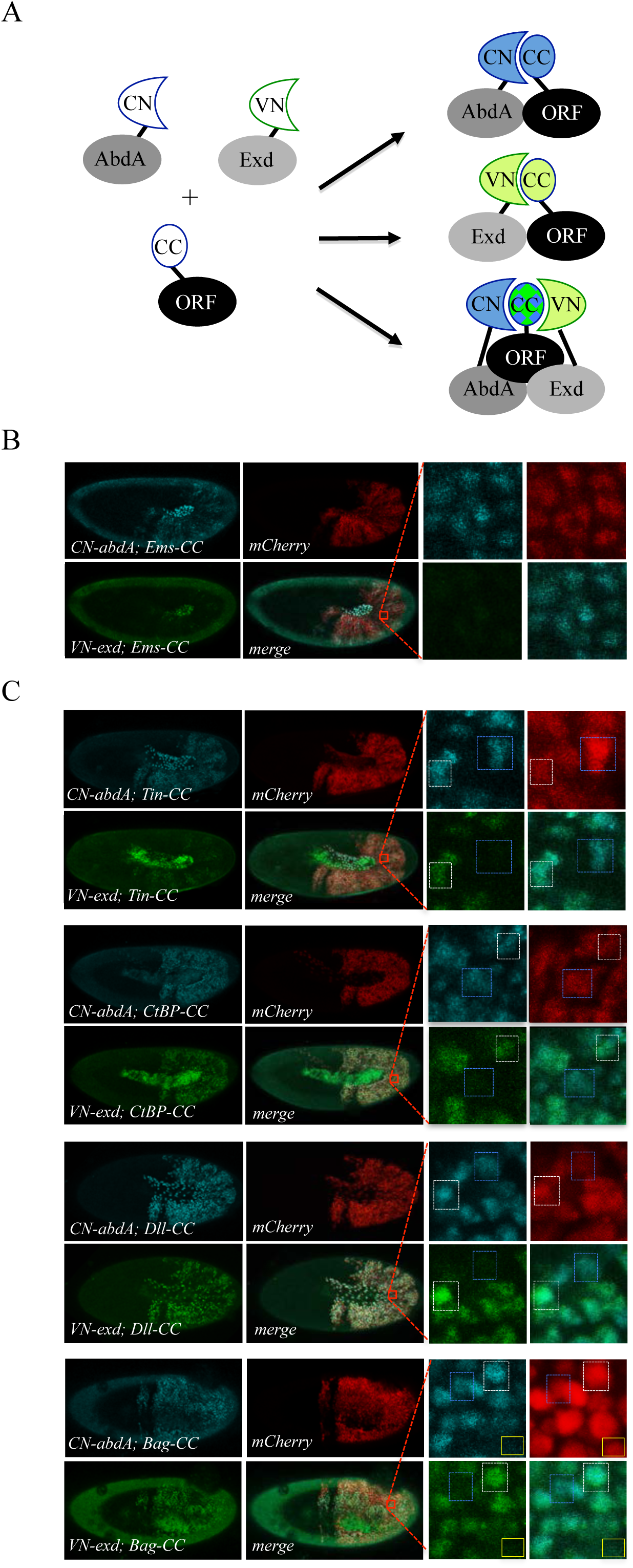
Using the multicolour BiFC library for analysing two different interactions in the same embryo. **A.** Principle of the bicolour BiFC. The AbdA and Extradenticle (Exd) cofactor are respectively fused to the CN or VN fragment, which can complement with the CC fragment of a co-expressed ORF when interaction occurs. The simultaneous expression of the three fusion proteins allows assessing Venus- and Cerulean-based BiFC in the same cell. Bicolour BiFC results from the interaction of the ORF-CC with both CN-AbdA and VN-Exd, therefore potentially in the context of AbdA/Exd complexes *in vivo*. **B.** Illustrative confocal capture of stage 10 embryo expressing Empty spiracles (Ems) fused to CC, together with AbdA and Exd fusion proteins, as indicated. BiFC is only occuring between AbdA and Ems, as expected from previous observation (see Table S8). **C.** Illustrative confocal captures of stage 10 embryos expressing CN-AbdA, VN-Exd and ORF-CC constructs, as indicated. Enlargements are provided for each embryo. White-dotted boxes depict nuclei where the ORF-CC interacts with both AbdA and Exd. Blue-dotted boxes depict nuclei where the ORF-CC interacts only with AbdA. Yellow-dotted boxes depict nuclei with absence of interaction. Fusion proteins are under the control of the *abdA-Gal4* driver. All expressing cells are recognized with the mCherry reporter. See also Supplementary Figures 13 and 14 and Table S8.

In order to identify such partners, we considered a set of 37 TFs fused to VN and previously described to be positive with VC-AbdA (^25^ and this work: Table S8). This set of TFs was tested with the VC-Exd fusion protein and BiFC was analysed in the epidermis of stage 10 embryos with the *abdA-Gal4* driver, as previously described ^46^. Results showed that 22/37 TFs were positive with Exd (Supplementary Figures 13 and 14 and Table S8), highlighting that the majority of AbdA-positive interactions was also positive with Exd, but also that many AbdA interactions might occur independently of the partnership with Exd *in vivo*.

We next selected four TFs (Tin, CtBP, Distalless (Dll) and Bagpipe (Bag)) that were positive with AbdA and Exd and present in the ORF-CC library to perform bicolour BiFC. An additional TF, Empty spiracles (Ems), which was positive with AbdA (^25^ and Figure 3B) but negative with Exd (Supplementary Figure 14), was also considered for the specificity control. The corresponding ORF-CC fusions were expressed together with CN-AbdA, VN-Exd and a mCherry reporter to trace Gal4-expressing cells in the embryo. Analysis with Ems-CC confirmed the respective positive or negative interaction status with AbdA or Exd, validating the specificity of BiFC upon the co-expression of the three fusion proteins in the same embryo (Figure 6B). Analysis with the four other TFs showed that the majority of expressing nuclei (as assessed with the mCherry reporter) were positive for both green and blue fluorescent signals, thus demonstrating that simultaneous bicolor BiFC was efficient with different types of TFs in the live *Drosophila* embryo (Figure 6C). Interestingly, a close-up in the embryo revealed that several nuclei were only positive in the Cerulean channel, indicating a specific positive interaction status with AbdA (blue-dotted boxes in Figure 6C). Green-only nuclei could not be found in the case of the four positive TFs, although Exd is expressed at the same level as AbdA. This suggests that BiFC with Exd is dependent on the concomitant interaction with AbdA and could therefore not be revealed outside AbdA/Exd complexes. Few red-positive nuclei were also negative for the two fluorescent signals (as illustrated with the yellow box in the case of Bag-CC in Figure 6C), highlighting that the interaction with both AbdA and Exd was actively inhibited in these specific nuclei.

Together, these observations confirm that the ORF-CC library is compatible for performing bicolour BiFC, providing a unique opportunity to reveal and investigate cell-specific regulatory mechanisms of two different PPIs *in vivo*.

Since the CC fragment is compatible for doing BiFC with the N-terminal fragment of Venus and Cerulean, we tested whether it could also complement with the N-terminal fragment of an additional GFP-derived protein called super-folder GFP (sfGFP). BiFC with sfGFP is described to rely on the complementation property between a long N-terminal fragment of 214 residues (sfGFPN) and a short C-terminal fragment of 19 residues (sfGFPC, ^47^). In addition to having distinct spectral properties from Venus and Cerulean (Figure 7A-A’), sfGFP has also a shorter maturation time 47.

**Figure 7.**
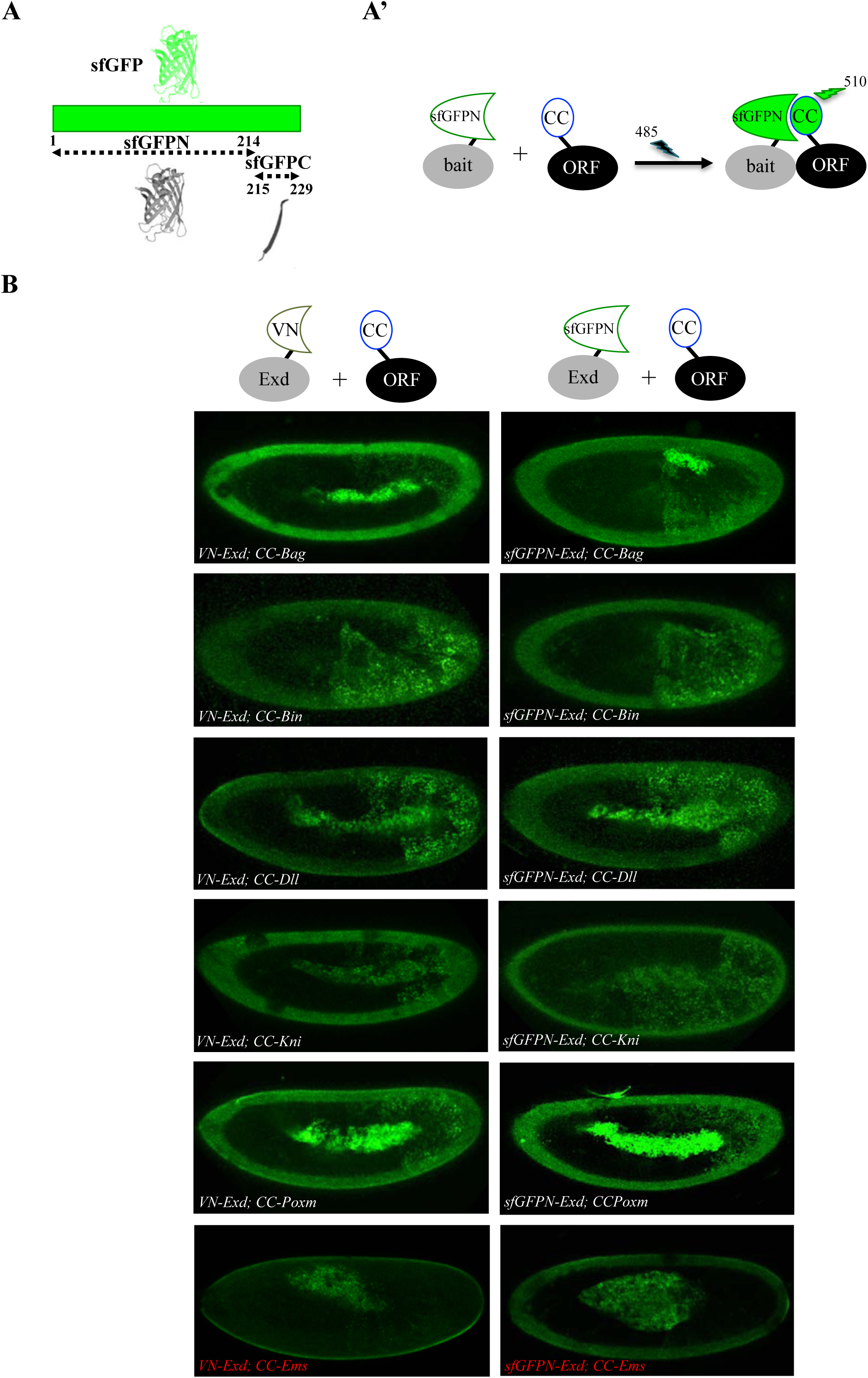
Adding the sfGFP to the fluorescence repertoire of the multicolour BiFC library. **A.** Split sfGFP fragments that are normally used for BiFC. **A’.** Principle of BiFC with the N-terminal fragment of sfGFP (sfGFPN) and the C-terminal fragment of Cerulean (CC). Excitation and emission wavelengths of the sfGFP are indicated. **B.** Illustrative confocal capture of BiFC obtained between ORF-CC constructs and VN-Exd (left panels) or sfGFPN-Exd (right panels), as indicated. Fusion proteins are expressed with the *abdA-Gal4* driver and BiFC is analyzed in the epidermis of stage 10 embryos. Absence of interaction with Empty Spiracles (Ems) is highlighted in red.

Here, we tested the complementation between sfGFPN and CC fragments by considering five TFs that were positive with Exd in our previous BiFC analyses. The sfGFPN fragment was fused at the N-terminus of Exd, making a sfGFPN-Exd fusion protein with the same fusion topology as previously done with the VN fragment (Materials and Methods). BiFC was analysed with excitation and emission wavelengths of the sfGFP. Results showed that the sfGFPN-Exd fusion protein could complement with the CC fragment in the case of all five tested TFs (Figure 7B). Moreover, fluorescent signals were of similar brightness when compared to signals obtained with VN-Exd (Figure 7B). Finally, since the sfGFPN and sfGFPC fragments are described as having a strong auto-affinity ^47^, we also performed BiFC with the negative control Ems-CC. No fluorescent signal was obtained between Ems-CC and sfGFPN-Exd (Figure 7B), confirming that the affinity between the sfGFPN and CC fragments is not strong enough to artificially induce complex formation between the two non-interacting proteins. Together these results establish yet another novel combination of fragment complementation for BiFC, adding to the fluorescence repertoire that can be used with the multicolor BiFC library.

### Using the multicolor BiFC library to visualize PPIs in the overlapping endogenous expression domains

The multicolor BiFC library is under the control of UAS sequences. Pilot tests were performed as previously described ^9,25^, using a unique Gal4 driver that reproduces the expression domain of the Hox protein in the embryo (Figure 8A). The same rationale could be applied with a Gal4 driver reproducing the expression profile of the ORF (Figure 8B). Under these conditions, BiFC signals are visualised in the expression domain of only one interacting partner. Assessing BiFC in a context where the two candidate partners are expressed in their endogenous domains could therefore improve the confidence in the interpretation.

**Figure 8.**
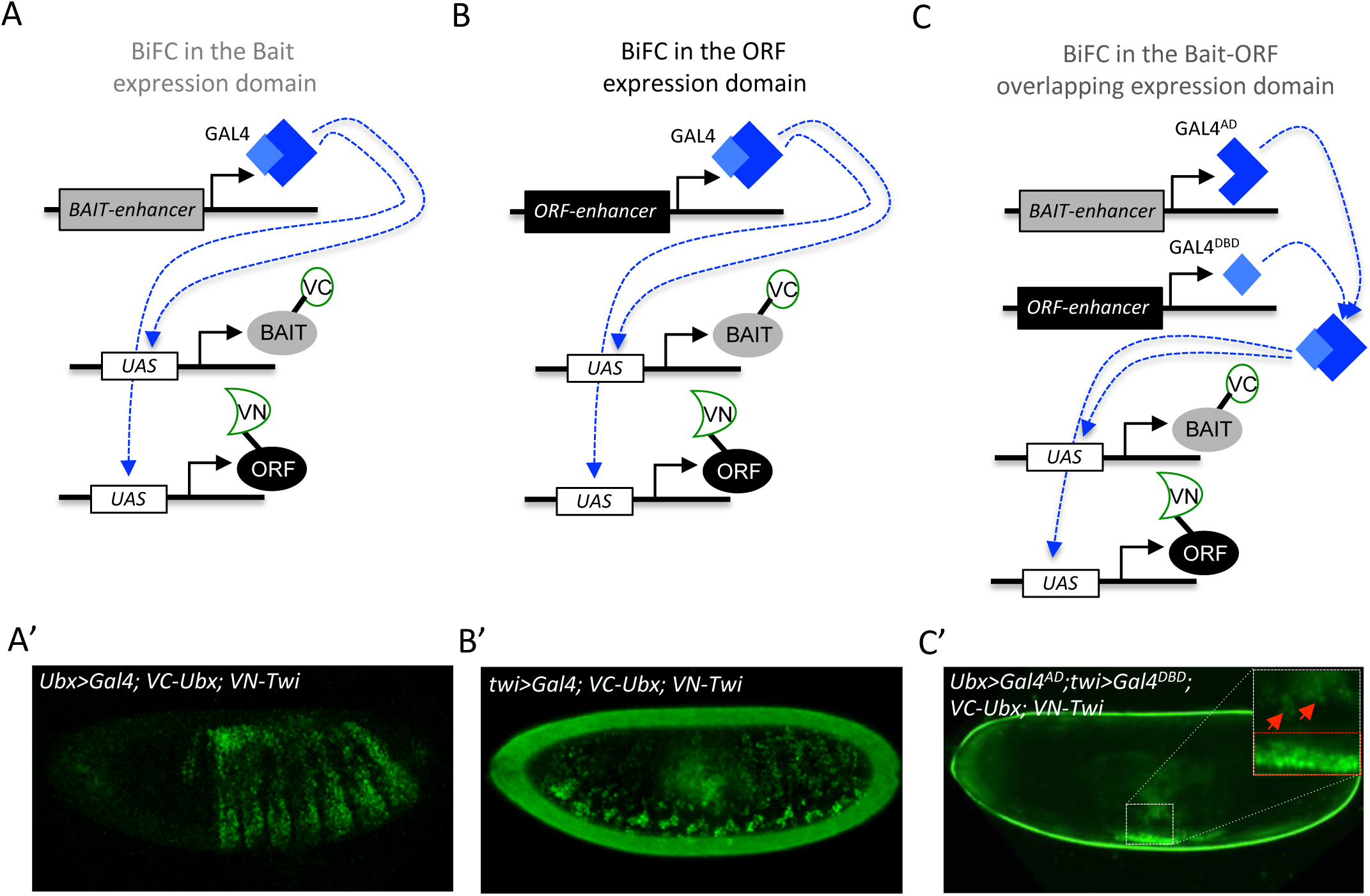
Coupling the multicolor BiFC library to the split-Gal4 system to visualize interactions in the overlapping expression domain of the two partners. **A.** Principle of BiFC with a unique Gal4 driver reproducing the expression profile of the bait protein (for example *Hox-Gal4* driver). **B.** Principle of BiFC with a Gal4 driver reproducing the expression profile of the ORF. **C.** Principle of BiFC upon the independent expression of Gal4 moieties (Gal4^AD^ and GAL ^DBD)^ by using two different enhancers from the bait- or ORF-encoding gene. This system allows producing a functional Gal4 protein in the overlapping expression domain of the two enhancers, therefore assessing BiFC in cells that normally express both the bait and the ORF. **A’-C’.** Illustrative confocal pictures of BiFC obtained upon the expression of Ultrabithorax (Ubx) and Twist (Twi) fusion proteins by using a Ubx-Gal4 (A’), twi-Gal4 (B’) or the split-Gal4 (Ubx-Gal4^AD^/twi-Gal4^DBD^: C’) system.

We have prospected towards this direction by considering the split-Gal4 system ^48^, which consists in reconstituting an active Gal4 protein upon association of two separate Gal4-DNA-binding (Gal4^DBD^) and Gal4-activation (Gal4^AD^) domains. This property can be used to activate any UAS-driven gene in cells that will independently co-express the Gal4^DBD^ and Gal4^AD^ moieties, for example under the control of two different enhancers ^48^. We reasoned that the split-Gal4 system could be used to assess BiFC specifically in the overlapping expression domain of the two candidate partners (Figure 8C). This strategy implies having enhancers that could reproduce the expression profile of each candidate partner and that are available for Gal4^DBD^ and/or Gal4^AD^ expression. A number of enhancers have been used for this purpose in the Janelia fly line collection (https://www.janelia.org/project-team/flylight). In addition, Mimic transposons have also been designed to be compatible with the split-Gal4 system ^49,50^. Altogether these genetic tools allow BiFC to reproduce the expression of thousands of genes in *Drosophila* using the split-Gal4 system.

Here the suitability of the split-Gal4 system with BiFC was tested by considering the interaction revealed between Ubx and the mesodermal TF Twist (Twi) ^25^. We used two enhancers from the Janelia collection that could recapitulate the expression profile of either *Ubx* or *twi* in the embryo. Each enhancer was first tested with the classical UAS/Gal4 system, leading to BiFC prominently visible in the epidermis or somatic mesoderm, respectively (Figure 8A’-B’). The same *Ubx* and *twi* enhancers were engineered with the split-Gal4 system, and were used to respectively express the Gal4^AD^ or Gal4^DBD^ moiety. In this genetic context, weak BiFC signals could be observed in few cells of the somatic mesoderm (red arrows in the enlargement of Figure 8C’), while strong BiFC signals was easily detected in the visceral mesoderm of the midgut (highlighted with the red-dotted box in the enlargement of Figure 8C’). This result confirms that the split-Gal4 system can be used with the multicolor BiFC library to analyze PPIs in the overlapping expression domain of the two interacting partners *in vivo*, thus reproducing more closely the endogenous interaction.

## Discussion

### A ready-to-use fly library for analyzing the interactions of hundreds of TFs *in vivo*

We present a new fly line library called multicolor BiFC library that currently allows using 453 TFs for testing PPIs *in vivo*. This library contains two different sets of fly lines. The first set contains 235 TFs fused to VN and derives from previous work ^9,10^ or from swapping experiments with the original FlyORF library. Although swapping experiments were performed for TF-encoding ORFs, it should be noticed that the FlyORF library actually covers around 3000 ORFs and is therefore not limited to TFs only. Proof of principles described with the VN-swapping fly lines could apply to many more ORFs since BiFC is compatible with different types of proteins. Moreover, using the complementary VC-swapping fly line ^23^ enables the full repertoire of the FlyORF library to be used for Venus-based BiFC in *Drosophila*.

The second set of the multicolor BiFC library consists of 326 ORF-CC constructs that are compatible with the first set of ORF-VN fly lines for doing Venus-based BiFC. ORF-CC fly lines are also compatible for doing Cerulean- or sfGFP-based BiFC with a CN- or sfGFPN-fused bait protein, respectively. This property allows multicolor BiFC experiments, which was here demonstrated with the analysis of interaction properties of different TFs in the context of the AbdA/Exd partnership. The multicolour BiFC system provides an unprecedented opportunity to dissect cell-specific regulatory mechanisms involving three interacting proteins *in vivo*.

### BiFC library enables high-confidence large-scale interaction screens

Among the 260 TFs that have been used for the large-scale interaction screen, 35 were tested in fusion with VN or CC to assess for reproducibility of BiFC when using a different complementation strategy (Table S9). This analysis showed that only 2/35 TFs (Hr83 and Ravus) did not reproduce the same result with VC-AbdA or VN-AbdA (Table S9). This observation highlights that the fusion topology is of minor incidence (6%) for causing false negatives, suggesting that PPIs are rarely completely abolished by inappropriate fusion topologies. In addition, it should be noted that the AbdA (and Ubx) fusion proteins used in this study were originally generated with an exaggerated short linker region (three residues long) to better measure the potential influence of fusion topologies when establishing BiFC in *Drosophila* ^9^. Using bait proteins fused to the Venus or Cerulean fragment with a more classical linker region (GGGSGG) could further diminish the influence of the fusion topology without increasing the risk of revealing indirect PPIs.

Our screen revealed that half of the 260 TFs could interact with Ubx or AbdA. As expected, a large proportion of interactions were common to the two Hox proteins, which reflects their ability to control a number of identical developmental processes during embryogenesis. The diversity of positive TFs reflects the likely high potential of Hox proteins to engage in many different types of interactions *in vivo*, as previously shown for Ubx ^51^. The screen was performed with a unique *Hox-Gal4* driver and therefore in conditions that do not reproduce the expression profile of the tested ORF. Despite this, we found that TFs that are endogenously more frequently co-expressed with Ubx and AbdA during embryogenesis were significantly enriched among the positive interactions.

Specificity of the BiFC screen was further demonstrated by the identification of several interactions that were exclusive to either Ubx or AbdA. Interestingly, Ubx- and AbdA-specific interactomes display opposite enrichment in molecular features involved in PPIs, including SLiMs and ordered/disordered regions. Moreover, the AbdA-specific interactome was significantly enriched in TFs that are expressed in tissues (gonad, heart) or cells (oenocytes) with known AbdA-specific functions. In addition, the experiments with HX-mutated AbdA and Ubx argue that many of these exclusive interactions are not merely false positive findings. Instead, the common and conserved HX motif is important for the integrity of each Hox-specific interactome.

Finally, the functional analysis of a selected pool of TFs expressed in the haltere disc showed that 80% of the interpretable phenotypes (22/30 TFs) were consistent with the interaction status found by BiFC in the embryo.

Altogether these results confirm that the multicolor BiFC library is suitable for performing large-scale interaction screens and revealing specific interactomes even for two closely related bait proteins.

### Suggestions for fine-tuning the specificity of the multicolor BiFC library screens

The multicolor BiFC library relies on the inducible UAS/Gal4 expression system, and caution should be taken regarding expression levels since high doses of protein expression could lead to artificial positive signals ^6^. In the case of a large-scale interaction screen with a unique Gal4 driver, we recommend using *P-Gal4* insertions corresponding to a mutant allele of the bait, as done here with the *Ubx-Gal4* or *abdA-Gal4* drivers. This genetic context allows expressing the bait protein at normal levels while eliminating one dose of endogenous competitive gene products. Specificity of BiFC could be validated by doing competition experiments (this work and ^25^). Alternatively, mutations affecting the interaction property of the bait protein could also be used as a specificity control, as shown here for several TFs with HX-mutated AbdA.

When possible, each candidate partner should also be expressed in its respective domain for assessing BiFC in the most relevant context. We showed that the split-Gal4 system could be used to visualize BiFC specifically in the overlapping expression domain of Ubx and Twi. The large number of fly lines compatible with the split-gal4 system makes this system very attractive for the future use of the multicolor BiFC library. Along the same line, the Janelia and Mimic collections propose hundreds of fly lines compatible with the LexAop/LexA-AD induction system. This expression system is also compatible with the multicolor BiFC library since UAS sequences of ORF-VN and ORF-CC constructs are swappable with LexAop sequences upon FLP/FRT-mediated recombination ^24^. The two candidate partners could then be expressed independently and in their respective domain by using the UAS/Gal4 system on the one side, and the LexAop/Lex-AD system on the other side. In this context, BiFC will only be analyzed in the overlapping expression domain of the two partners, as described with the split-Gal4 system. Since the ORF-CC library is inserted on the second chromosome, we have generated another LexAop-swapping fly line that is compatible for FRT-mediated recombination with this chromosome (see Materials and Methods).

In conclusion, the versatile inducible expression systems for thousands of enhancers in *Drosophila* and the multicolor BiFC library enable selection of biologically relevant conditions that better recapitulate the endogenous expression profile of each candidate partner, thus maximizing specificity and providing an unprecedented combination of possibilities for large-scale and/or in depth analysis of PPIs in a live animal organism.

## Materials and Methods

### Plasmid constructions

#### 1. Cloning of pTSVNm9short.attB

The FRT2-VNm9short fragment was PCR-amplified from plasmid pTSVNm9.attB with forward primer FRT2-VN-F (5’-TATGGATCCGAAGTTCCTATTCTCTACTTAGTATAGGAACTTCGATGGTGAGCAAGGGCG-3’) and reverse primer tub-R2 (5’-ACACTGATTTCGACGGTTACC-3’), thereby eliminating a stretch of 12aa between FRT2 and VNm9 and introducing flanking restriction sites BamHI and NotI. Plasmid pTSeGFP.attB (without the *yellow* marker insert) was digested with BamHI-NotI and the shortened FRT2-VNm9short fragment was inserted. Finally, the yellow marker gene was inserted via the XhoI site downstream of the fragment, resulting in pTSVNM9short.attB (for additional cloning details see ^23^).

#### 2. Cloning of pTSCN.attB

The N-terminal part of Cerulean (CN) including the FRT2 site was PCR-amplified from the Cerulean cDNA ^14,9^ with forward primer FRT2-VN-F (5’-TATGGATCCGAAGTTCCTATTCTCTACTTAGTATAGGAACTTCGATGGTGAGCAAGGGCG-3’) and reverse primer Not-CN-R (5’-ATAGCGGCCGCctaGGTGATATAGACGTTGTCG-3’), thereby introducing BamHI and NotI sites. Note, the start sequence of the N-terminal end of Venus and Cerulean is identical. Subsequently, the CN fragment and the yellow marker were introduced into pTSeGFP.attB the same way as described above, generating plasmid pTSCN.attB.

#### 3. Cloning pGW-CC.attB

To generate this plasmid, the 3-HA tag from pGW-HA.attB ^23^ was replaced with the C-terminal portion of Cerulean (CC). In brief, fragment CC was PCR-amplified from the Cerulean cDNA with primers CC-F (5’-ATAGGTACCTGCCGACAAGCAGAAGAACG-3’) and CC-R (5’-TATGCTAGCTTACTTGTACAGCTCGTCCATGCCG-3’). Note, in this construct no FRT2 sequence was included in front of the Cerulean fragment, as no swapping experiments are intended with the subsequent fly lines resulting from this construct. The CC fragment was inserted into pGW-HA.attB with KpnI-NheI digestion, thus releasing FRT2 fragment and 3xHA tag and generating plasmid pGW-CC.attB.

#### 4. Creating the TF-ORF-CC plasmid library

A detailed description of the involved Gateway cloning steps to generate such a library can be found in previous publications ^23,24^. In brief, the *Drosophila* transcription factor collection in the pDONR221 vector ^52^ was used as a source for the ORFs. From this collection the TFs were shutteled (Gateway subcloned) into the *Drosophila* expression vector pGW-CC.attB, thus fusing the ORF to the CC tag and equipping them with UAS regulatory promoter elements.

#### 5. Cloning of sfGFPN-Exd in pUASTattB

The N-terminal fragment of sfGFP (sfGFPN: residues 1-214 ^47^) was PCR amplified from the sfGFP cDNA and cloned in place of VN in the original VN-Exd construct in pUASTattB between EcoRI and XhoI restriction sites ^9^.

### Generation of fly strains

For all germline transformation experiments we used the ΦC31 integrase-mediated site-specific integration method.

#### 1. Generation of individual fly strains for promoter or tag swapping

*TSVNm9short-86Fb, TSCN-86Fb* strains: The constructs pTSVNM9short.attB and pTSCN.attB were injected into line *ΦX-86Fb.* Transgenic offspring was made homozygous for these transgenes and combined with an X chromosome-linked hsp70-flp construct. These strains provide either the Venus N-terminal tag or the Cerulean N-terminal tag for a swapping event at the 3’end of any UAS-ORF strain from FlyORF that is based on cloning with pGW-HA.attB.

For the *PSlexO-21F* strain the pPSlexO.attB plasmid was injected into line *ΦX-21F*, again followed by balancing the strain for the transgene and providing the hsp70-flp construct on the X-chromosome. This strain can be used to exchange the UAS-regulated promotor of the TF-ORF-CC library (at location 21F) for a lexA operator (lexO) promotor ^23,53^.

#### 2. Creating a transcription factor Cerulean tagged fly library (UAS-ORF-CC) on the second chromosome

The individual UAS-ORF-CC plasmids were combined in small pools and injected into *ΦX21F*. Transgenes that were not recovered after outcrossing and PCR identification were re-pooled and injected again. The PCR identification for this library was done by single-fly PCR and Sanger sequencing into the 5’ region of the genes. Individual stocks were created with balancer line (*y^−^w^−^; Bc gla / SM6a*). Pool injection, transgene identification and stock generation were described in detail previously ^23,24^.

#### 3. Swapping procedures: exchange of 3xHA tag for VNm9 or VNm9short

The presence of the mutated FRT2 site downstream of the ORF was used to replace the 3xHA tags of FlyORF lines for the Venus tags VNm9 or VNm9short. In brief, the ORF-3xHA lines were crossed to flies carrying a swap construct together with the hsp-flp transgene. After heat-shock treatment and outcrosses the swapping events (FLP/FRT-mediated in vivo events) can easily be tracked by screening for a specific marker combination, i.e. *y+w+* for a C-terminal tag exchange. This procedure was described in detail previously ^23,24^.

### BiFC analysis

Fly crosses, embryo preparation and BiFC observation in live embryos were performed as previously described ^9,10^. Briefly, observations were performed at least twice or three times (from two different night egg laying periods) with wild type or mutated Hox proteins, respectively. BiFC signals with wild type Hox proteins were considered as positive when the intensity was above the fluorescent background and reproducible in the expected proportion of embryos from independent preparations (around 200 embryos were mounted in each case for confocal observation). Fluorescence intensities are strongly fluctuating from one TF to another, which could be due to the influence of the fusion topology and/or a real variation in interaction affinity with the Hox protein. Identical parameters of confocal acquisition were applied with mutated Hox proteins or in competition experiments.

### Fly stocks

The *Ubx-Gal4* and *abdA-Gal4* lines were previously used ^9^. UAS-ORF-HA fly lines used for swapping and competition experiments are from the FlyORF library ^23^. *Ubx-Gal4^AD^* and *twi-Gal4^DBD^* are from Janelia (Bloomington stock numbers 70646 and 68953 respectively). RNAi against TFs are from Bloomington and were expressed with the *MS1096* driver combined or not with UAS-dicer in individuals heterozygous for the *abxpbxbx* mutation, which affects *Ubx* expression in the haltere disc ^41^. The transgenic BiFC library lines will be available from FlyORF (http://www.flyorf.ch).

### Statistical analyses

Wilcoxon statistical tests were performed by using Python and R in-house scripts. Statistical analysis of intrinsically disordered/ordered regions in Ubx- and AbdA-specific interactome was assessed by using the DisEMBL pipeline ^54^, with a 0.5 cut-off score in the COILS prediction for each residue (>= 0.5 is considered disordered). The SLiM enriched regions were predicted using SLiMPred ^55^, with a 0.1 cut-off score (>= 0.1 is considered as SLiM). Each residue was sorted into one of four categories using two tags (ordered/disordered and SLiM/non-SLiM). For each interactome, the number of residues with the same tags was divided by the total number of residues in that interactome, thus yielding four proportions that were compared across interactomes using a two-tailed two-proportions z-test (allowing correction for different sample sizes).

## Supplementary Figure legends

**Supplementary Figure 1.**
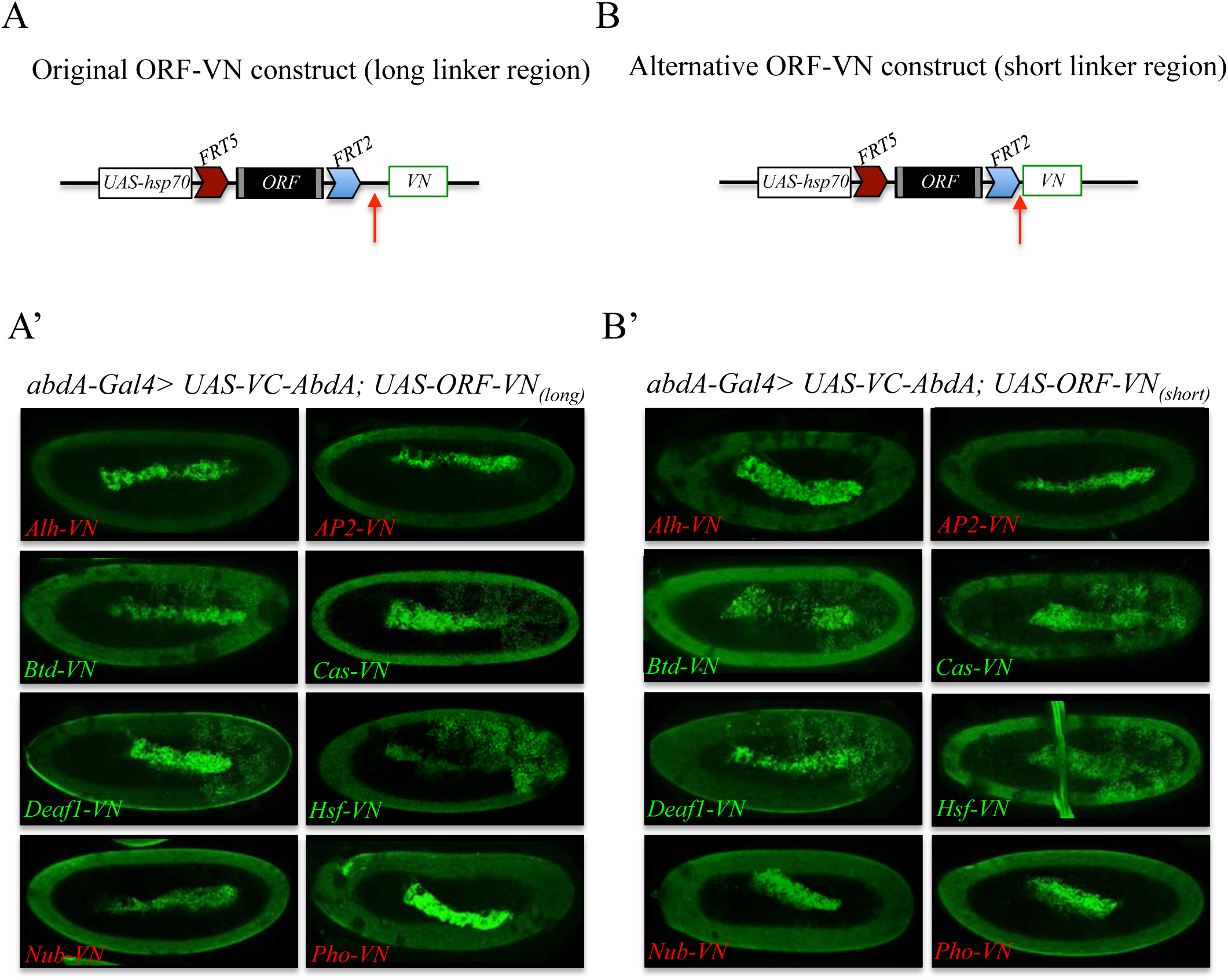
Comparison of Venus-based BiFC when using ORFs swapped with the original VN or VN-short tag line. **A.** Schematic representation of an ORF fused to VN with the original linker region (red arrow). **A’.** Illustrative confocal captures of BiFC resulting from the co-expression of ORFs fused to VN with the original linker region and the Hox protein AbdominalA (AbdA) fused to the complementary VC fragment. **B.** Schematic representation of an ORF fused to VN with the new short linker region (red arrow). **B’.** Illustrative confocal captures of BiFC resulting from the co-expression of ORFs fused to VN with the new short linker region and the Hox protein AbdominalA (AbdA) fused to the complementary VC fragment. Fusion proteins are expressed with the *abdA-Gal4* driver and BiFC is shown in the epidermis of stage 10 embryos. Note that the amnioserosa inside the embryo displays prominent autofluorescence. ORFs that are negative with AbdA are highlighted in red.

**Supplementary Figure 2.**
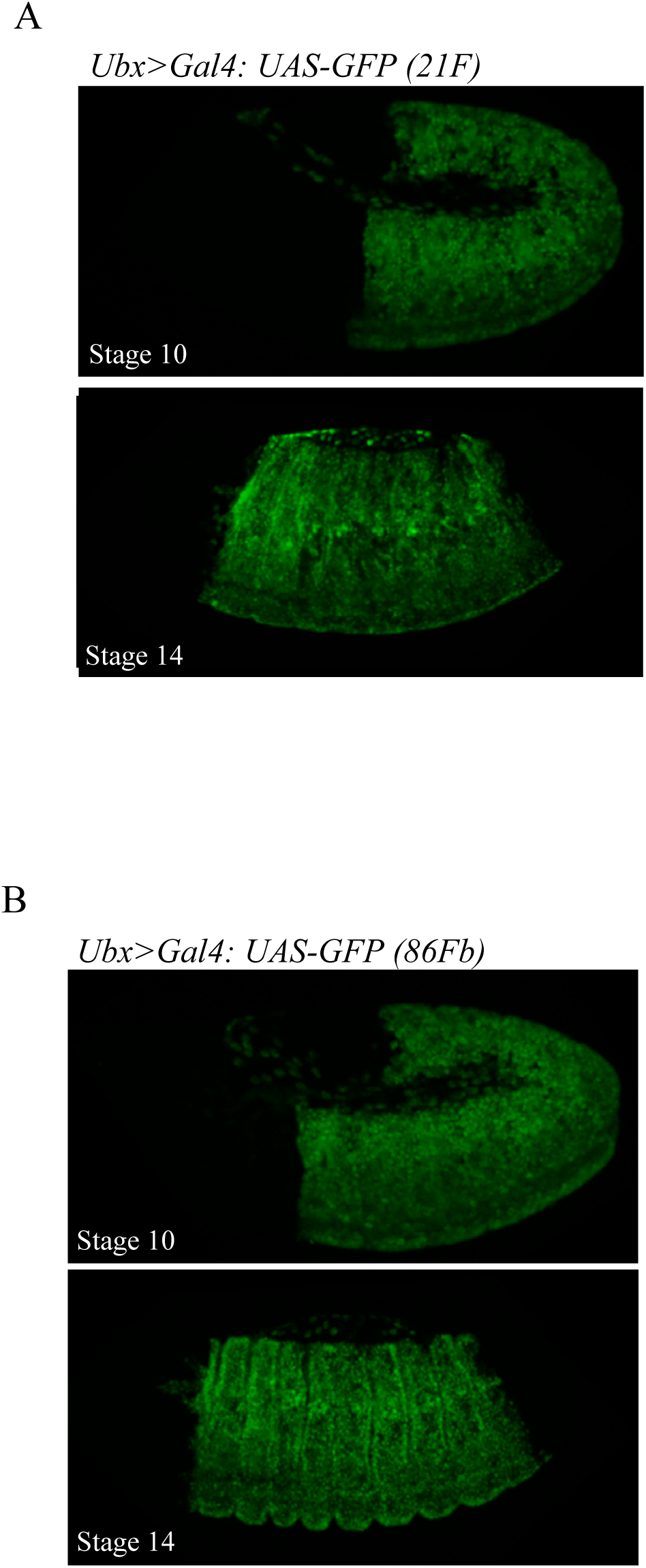
The 21F and 86Fb attP sites lead to similar expression levels. Illustrative confocal capture of UAS-driven GFP inserted in the 21F (A) or 86Fb (B) attP sites with the *Ubx-Gal4* driver. Two different stages are shown.

**Supplementary Figure 3.**
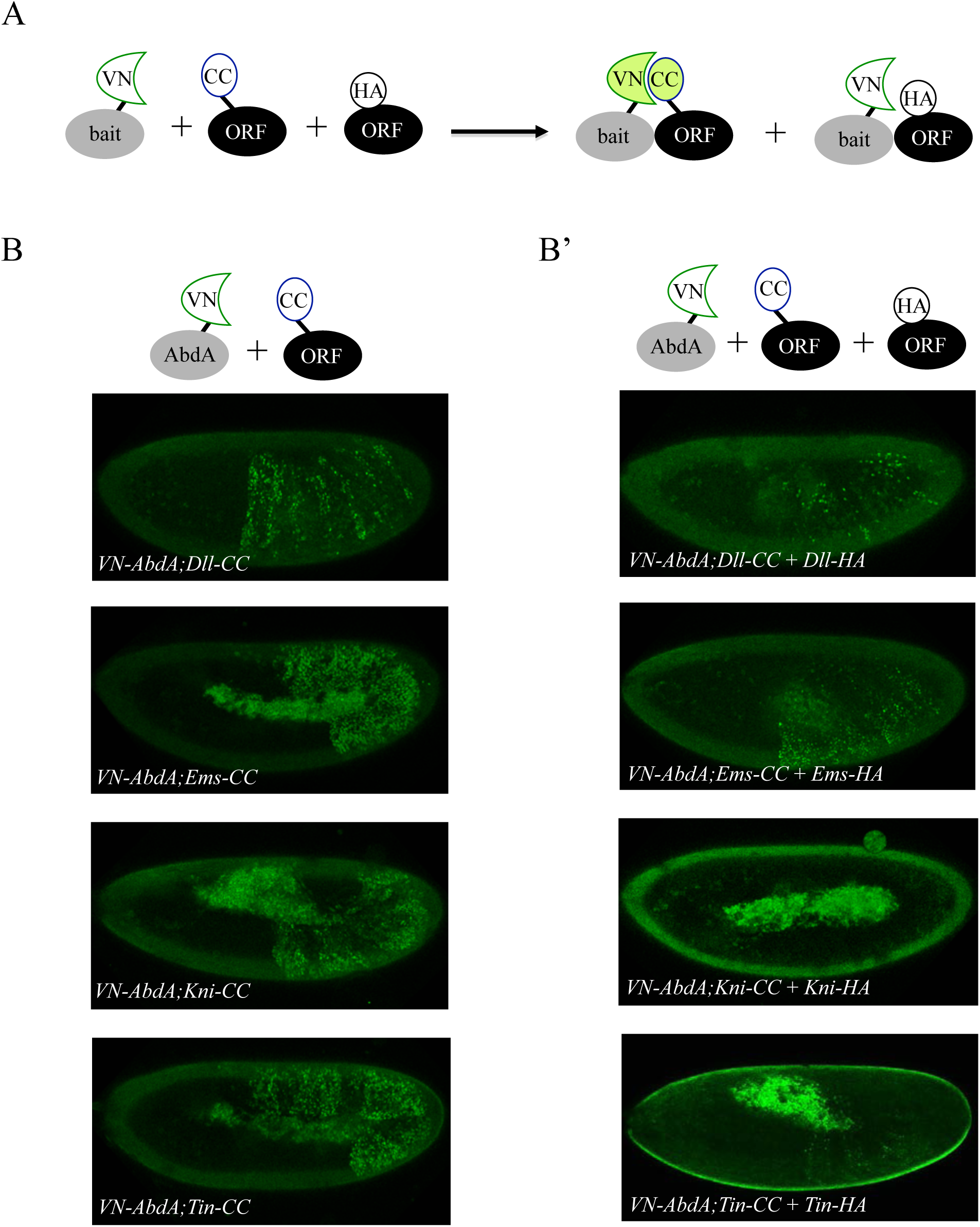
Co-expression of non-complementing ORF-3xHA can compete against BiFC signals obtained from ORF-CC and VN-AbdA constructs. **A.** Principle of the BiFC competition experiment with co-expression of a non-complementing HA-tagged ORF. **B.** Illustrative confocal captures of BiFC between VN-AbdA and the different ORF-CC constructs, as indicated. **B’.** Illustrative confocal captures of BiFC between VN-AbdA and the same ORF-CC constructs in the presence of the corresponding ORF-HA. Constructs are expressed with the *abdA-Gal4* driver and BiFC is analysed in the epidermis of stage 10/11 embryos. The co-expression of non-complementing ORF-HA strongly affects BiFC in all cases.

**Supplementary Figure 4.**
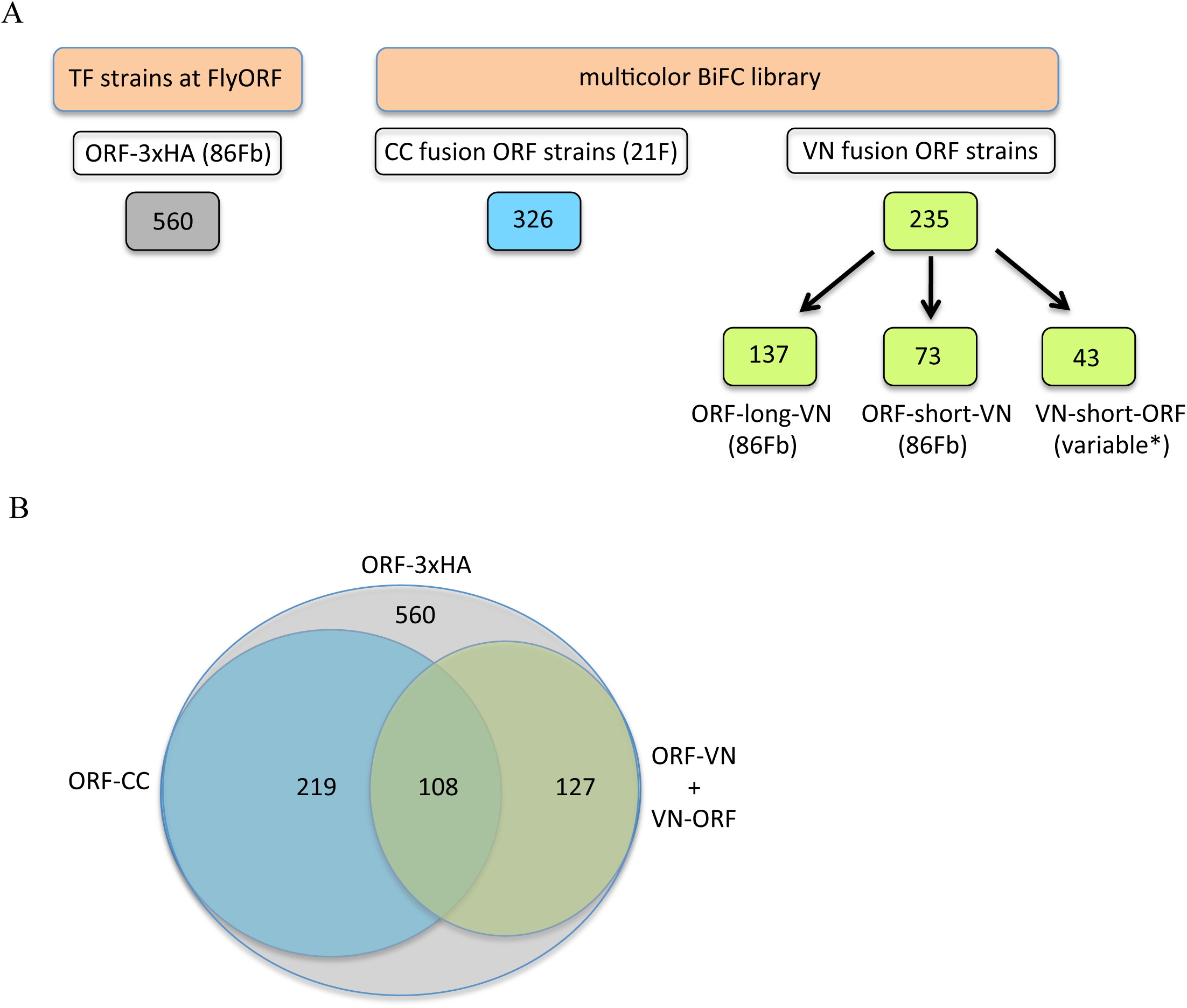
The TF-3xHA and multicolour BiFC libraries. **A.** Number of TF-3xHA and multicolour BiFC fly strains at FlyORF. Various insertion sites were used for the VN-short-ORF constructs ^10^. Some of the 234 TF strains exist in more than one VN version (see also Table S1). **B.** Repartition of the multicolour BiFC fly lines compared to the TF-3xHA library.

**Supplementary Figures 5a and 5b.**
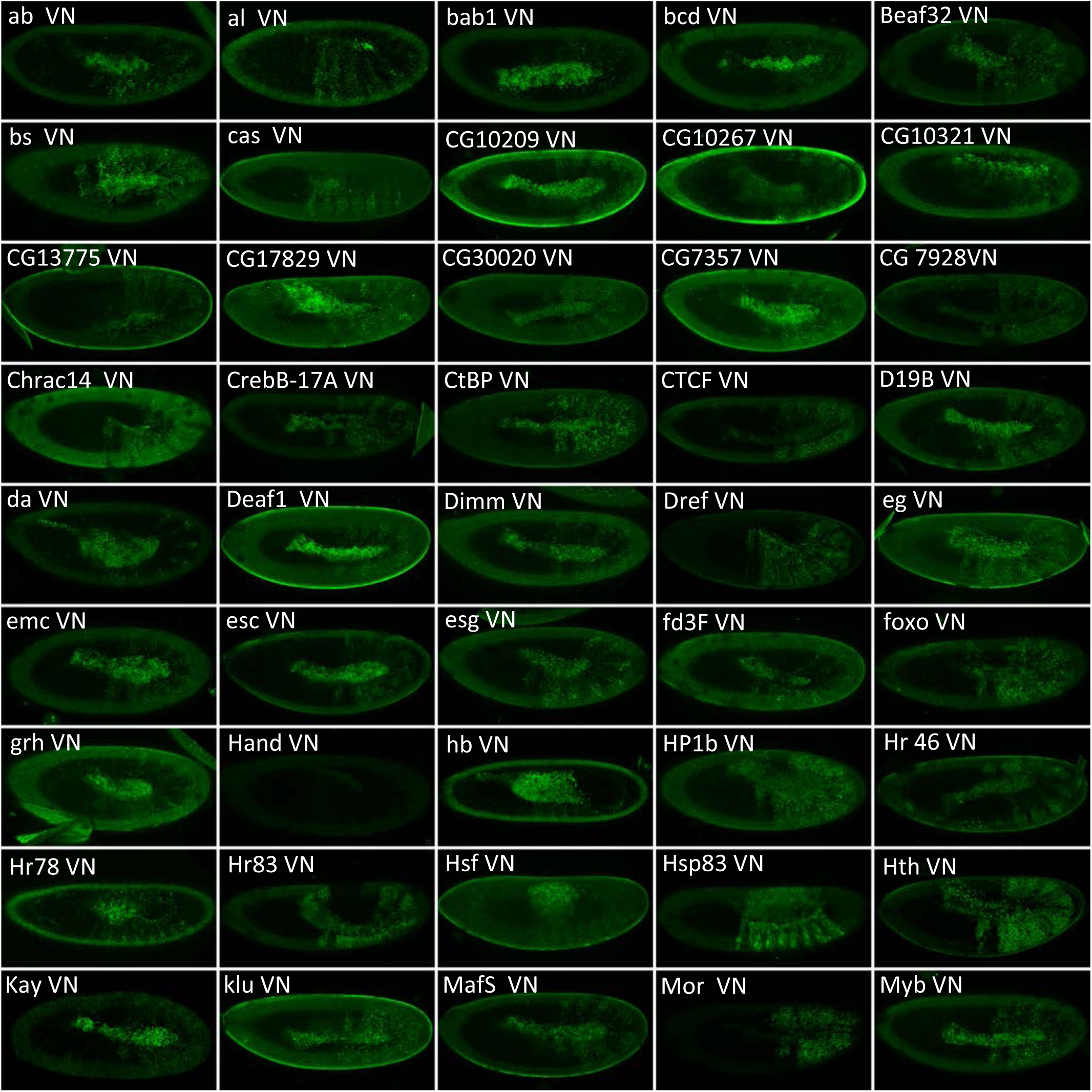
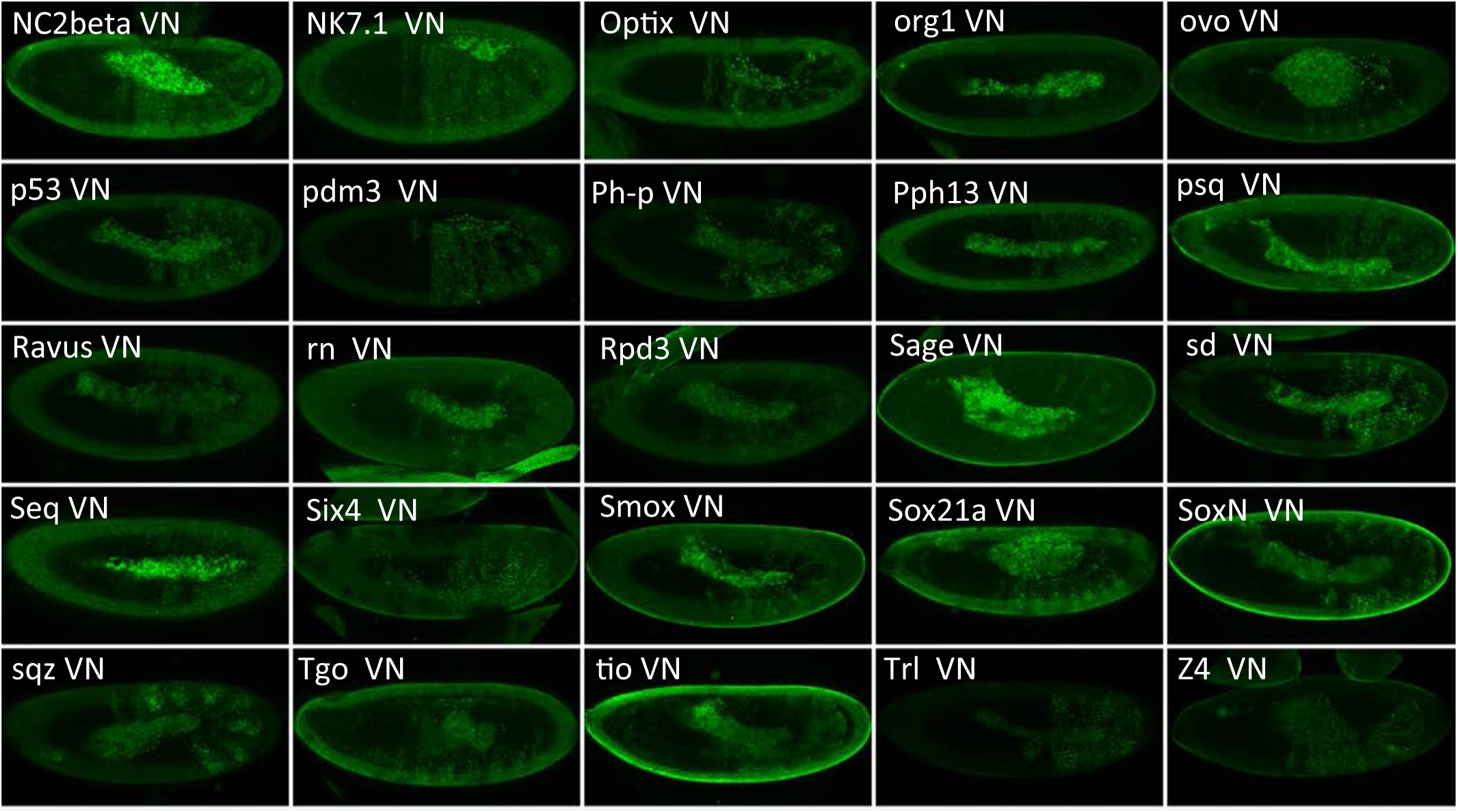
Illustrative confocal captures of Venus-based BiFC between ORF-VN and VC-Ubx, as indicated. Fusion proteins are expressed with the *Ubx-Gal4* driver and BiFC analysed in the epidermis of stage 10 embryos.

**Supplementary Figures 6a and 6b.**
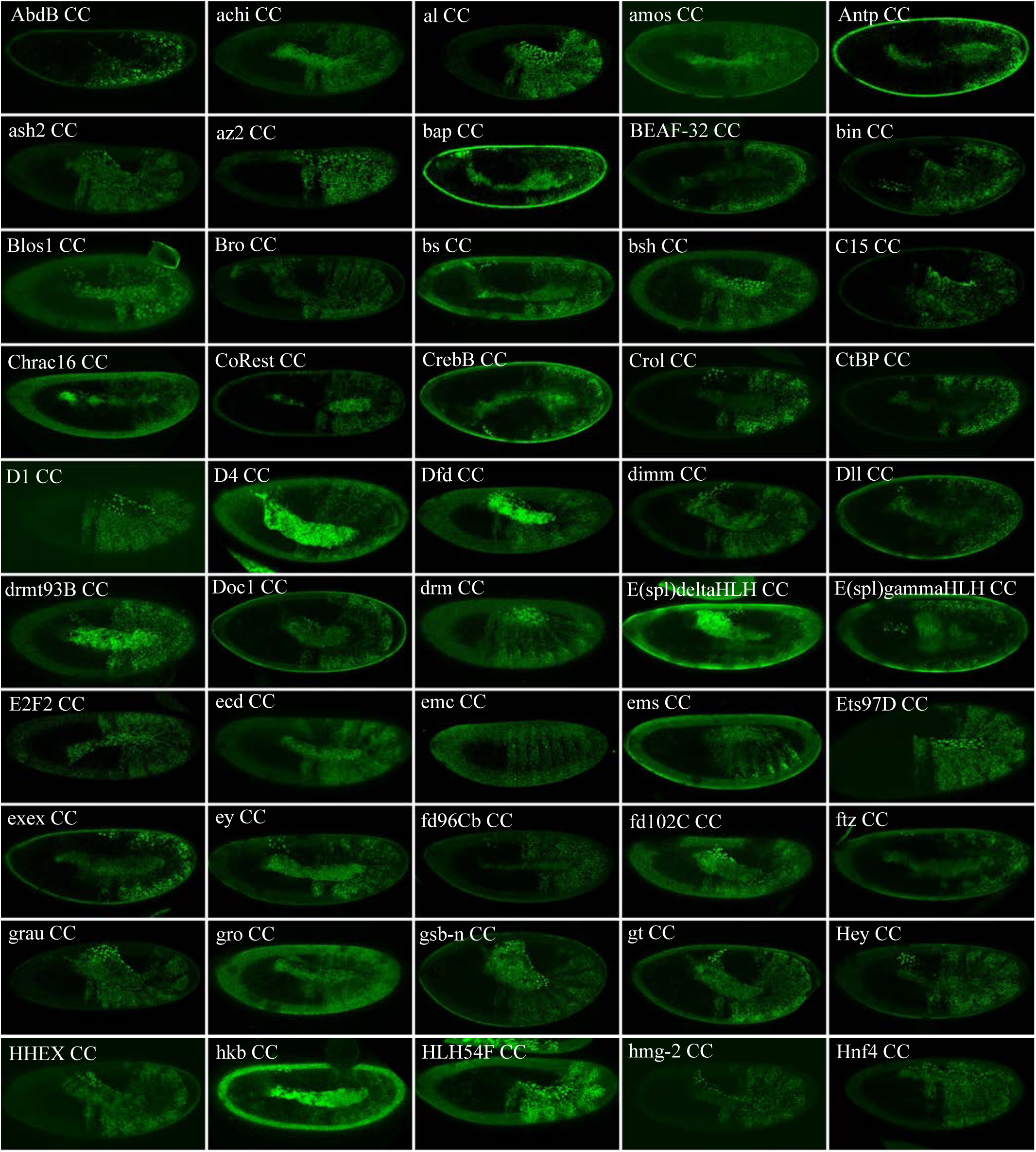
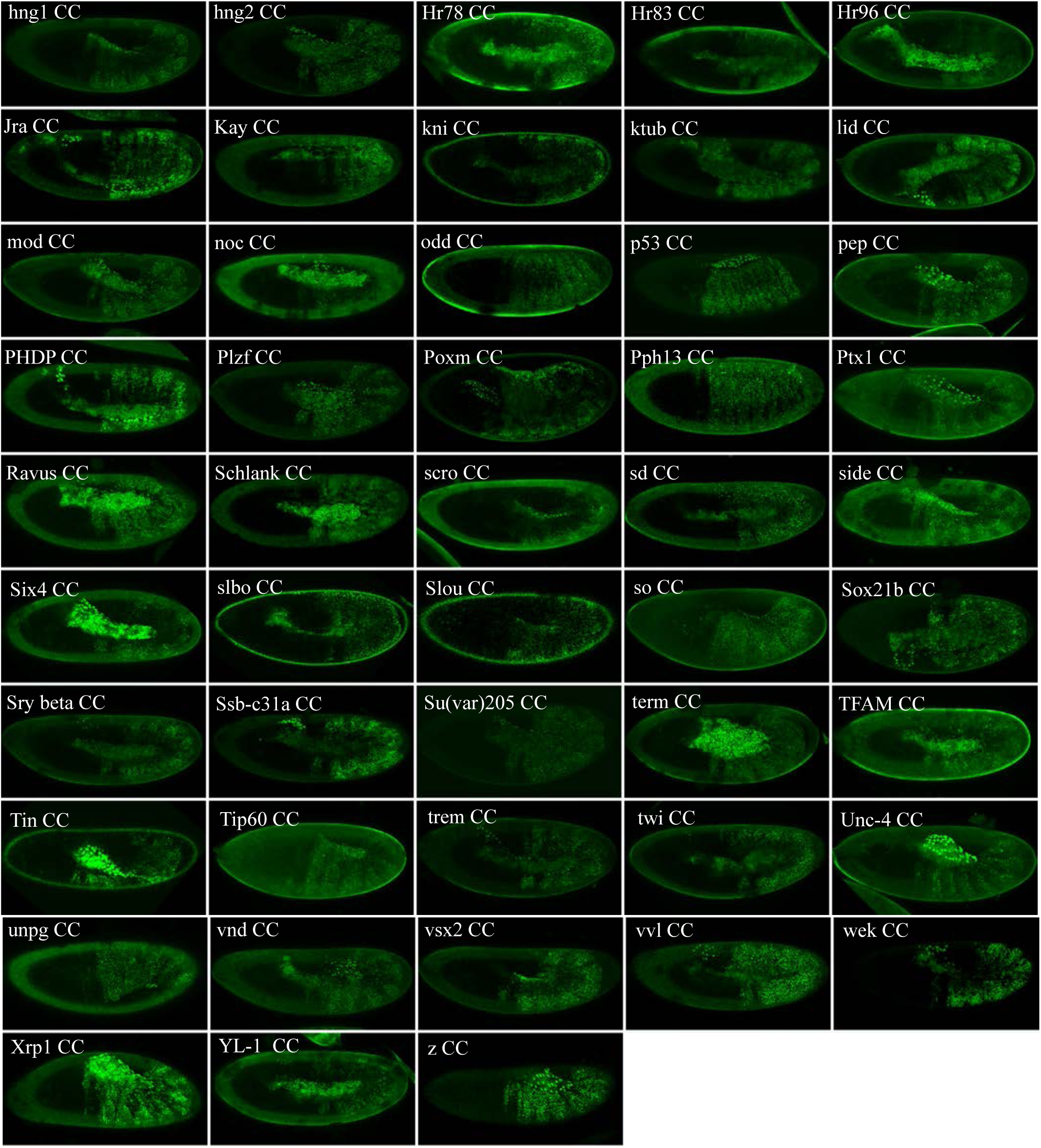
Illustrative confocal captures of Venus-based BiFC between ORF-CC and VN-Ubx, as indicated. Fusion proteins are expressed with the *Ubx-Gal4* driver and BiFC analysed in the epidermis of stage 10 embryos.

**Supplementary Figures 7a and 7b.**
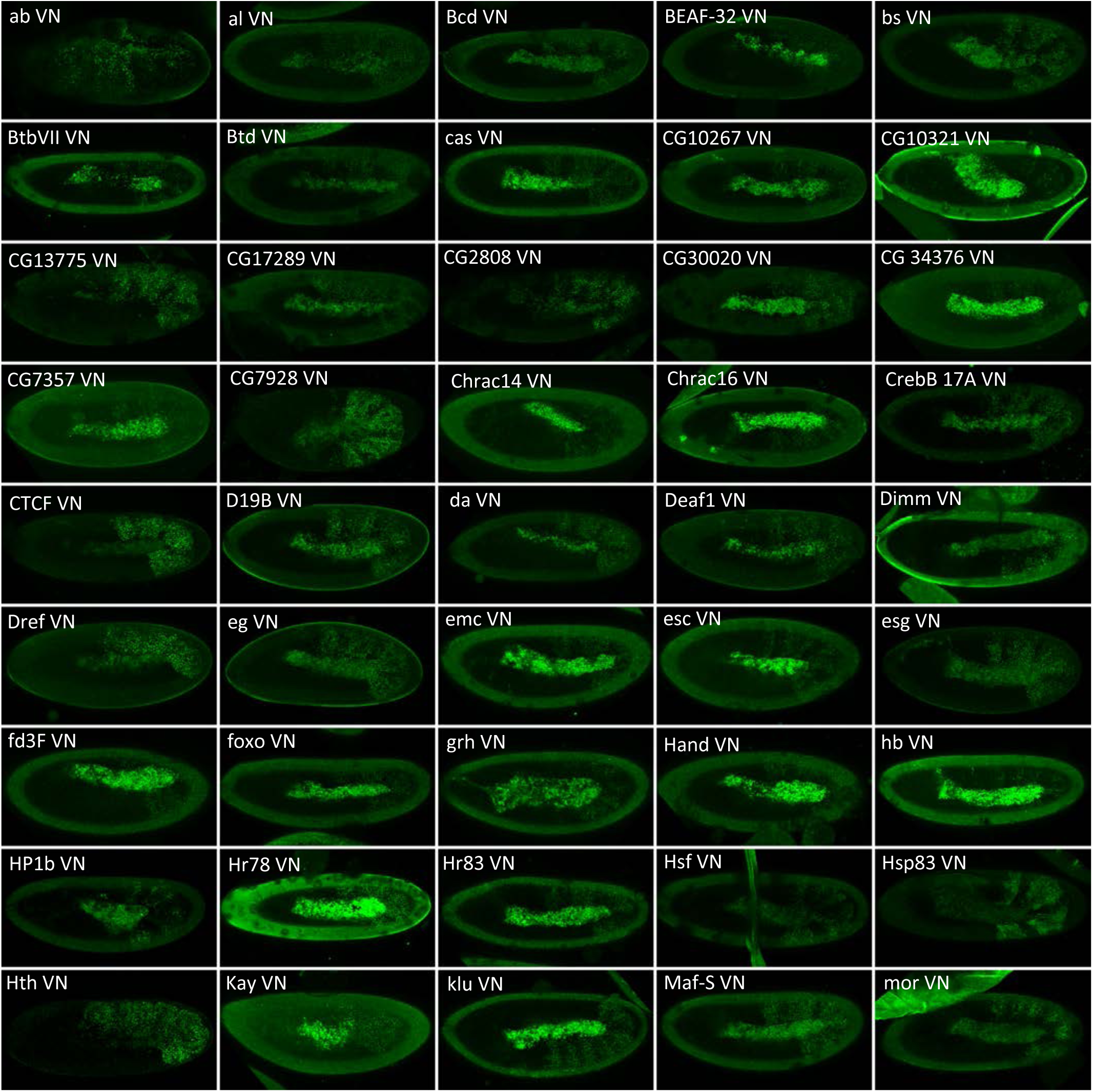
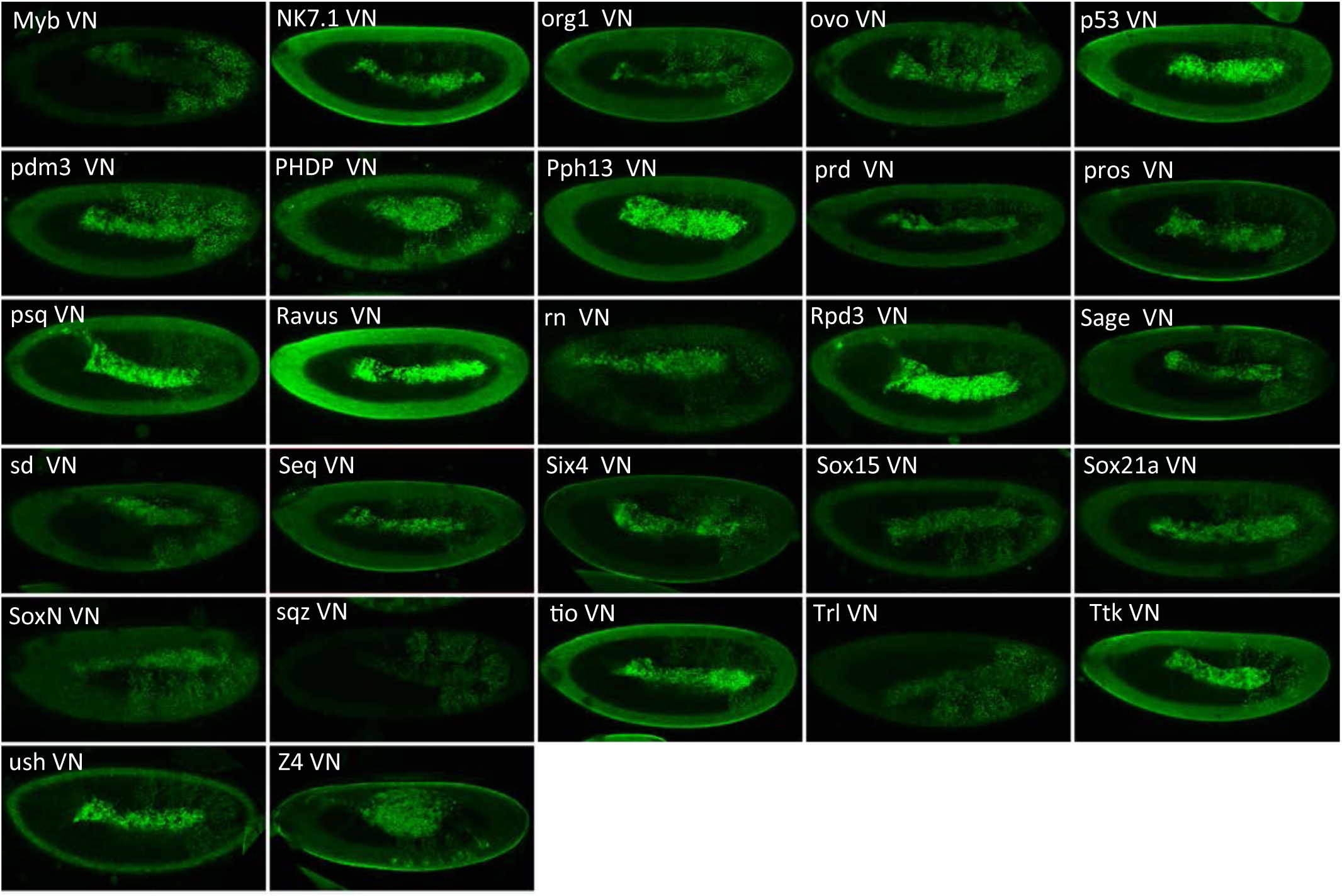
Illustrative confocal captures of Venus-based BiFC between ORF-VN and VC-AbdA, as indicated. Fusion proteins are expressed with the *abdA-Gal4* driver and BiFC analysed in the epidermis of stage 10 embryos.

**Supplementary Figure 8.**
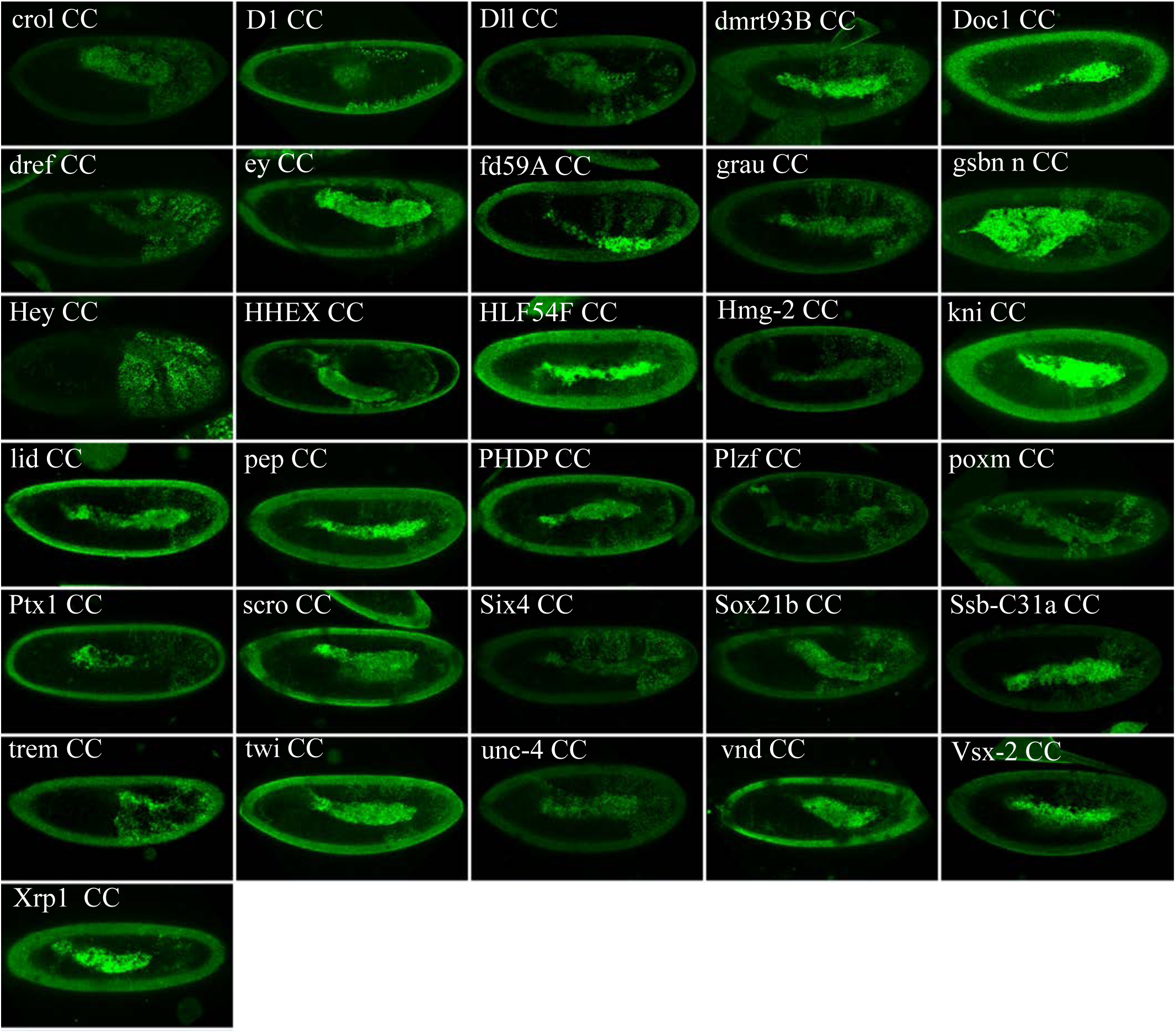
Illustrative confocal captures of Venus-based BiFC between ORF-CC and VN-AbdA, as indicated. Fusion proteins are expressed with the *abdA-Gal4* driver and BiFC analysed in the epidermis of stage 10 embryos.

**Supplementary Figure 9.**
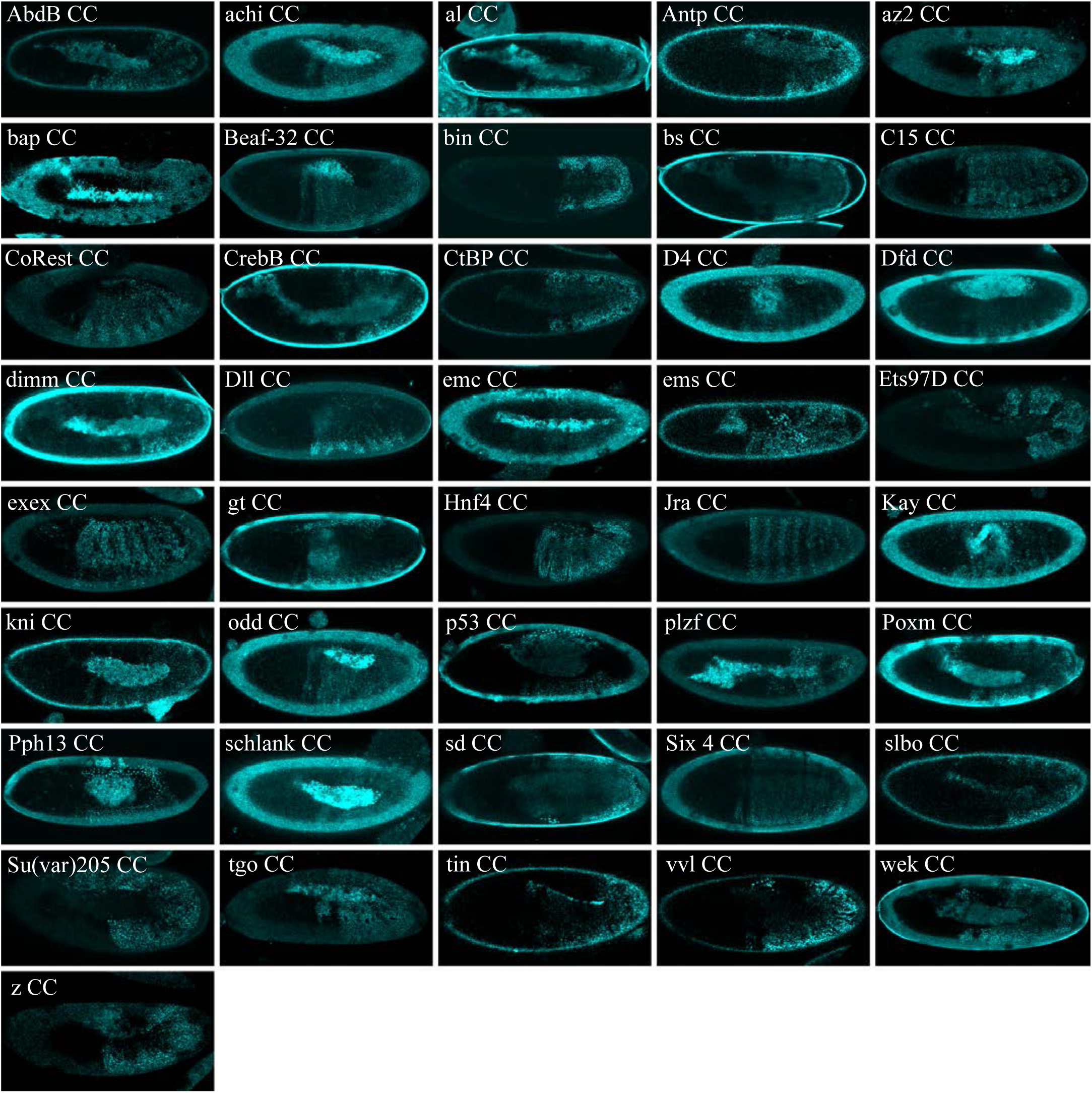
Illustrative confocal captures of Cerulean-based BiFC between ORF-CC and CN-AbdA, as indicated. Fusion proteins are expressed with the *abdA-Gal4* driver and BiFC analysed in the epidermis of stage 10 embryos.

**Supplementary Figure 10.**
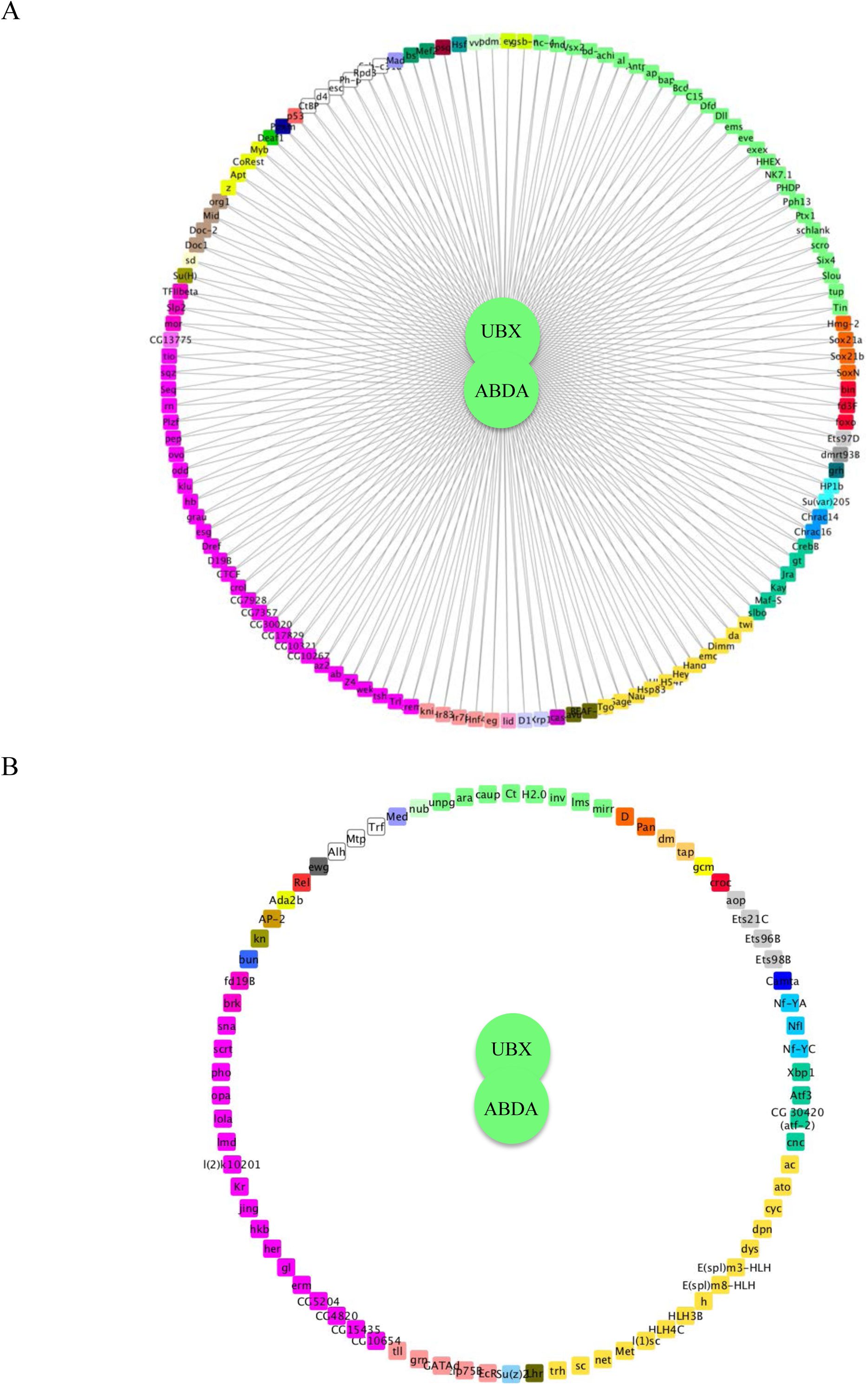
Representation of the common interactome (A) and negatome (B) between Ubx and AbdA. Colour code is as in Figure 3.

**Supplementary Figure 11.**
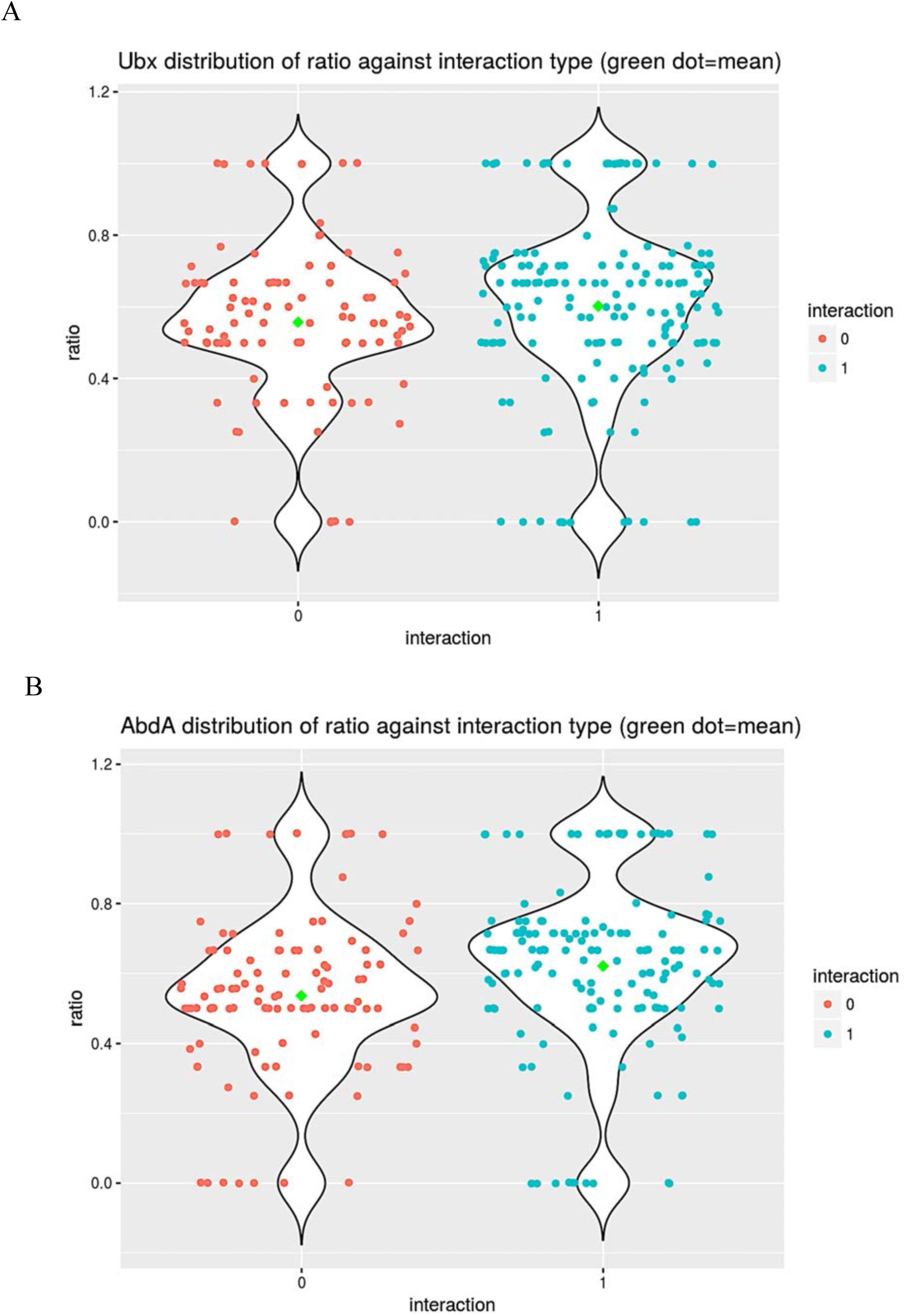
Analysis of the co-expression and interaction status of Ubx (A) or AbdA (B) and the positive ORFs. The distributions of the values taken by the ratio between the number of tissues in which the TF and Ubx or AbdA are co-expressed and the total number of tissues composing the TF expression domain during embryogenesis (among the 25 analysed developmental contexts) with regard to their interaction status (red and blue dots indicate negative (0) and positive (1) interaction status between the TFs and Hox proteins, respectively). Green circles show the means of the distributions.

**Supplementary Figure 12.**
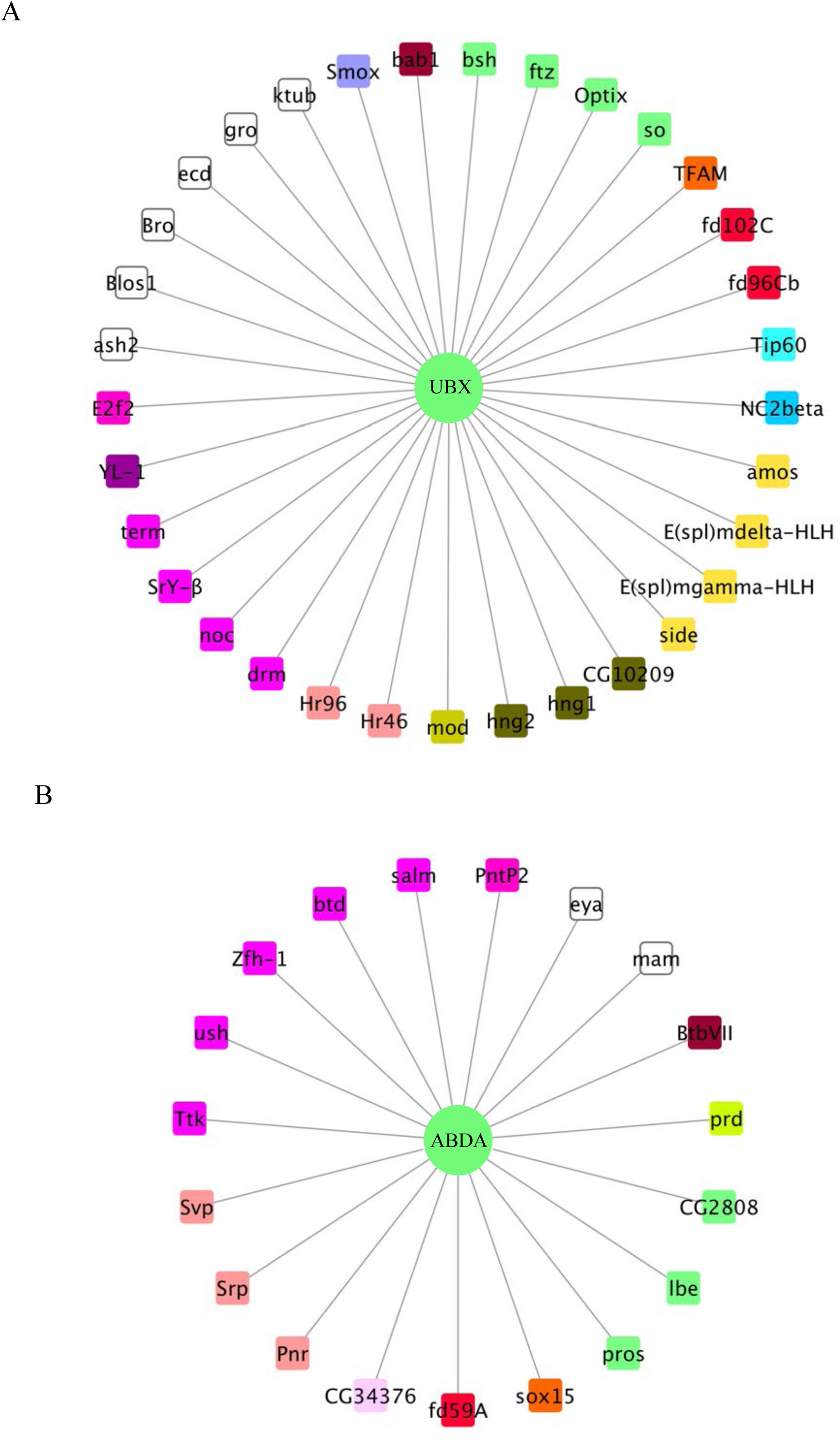
Representation of the Ubx- (A) and AbdA-specific (B) interactomes. Colour code is as in Figure 3.

**Supplementary Figure 13.**
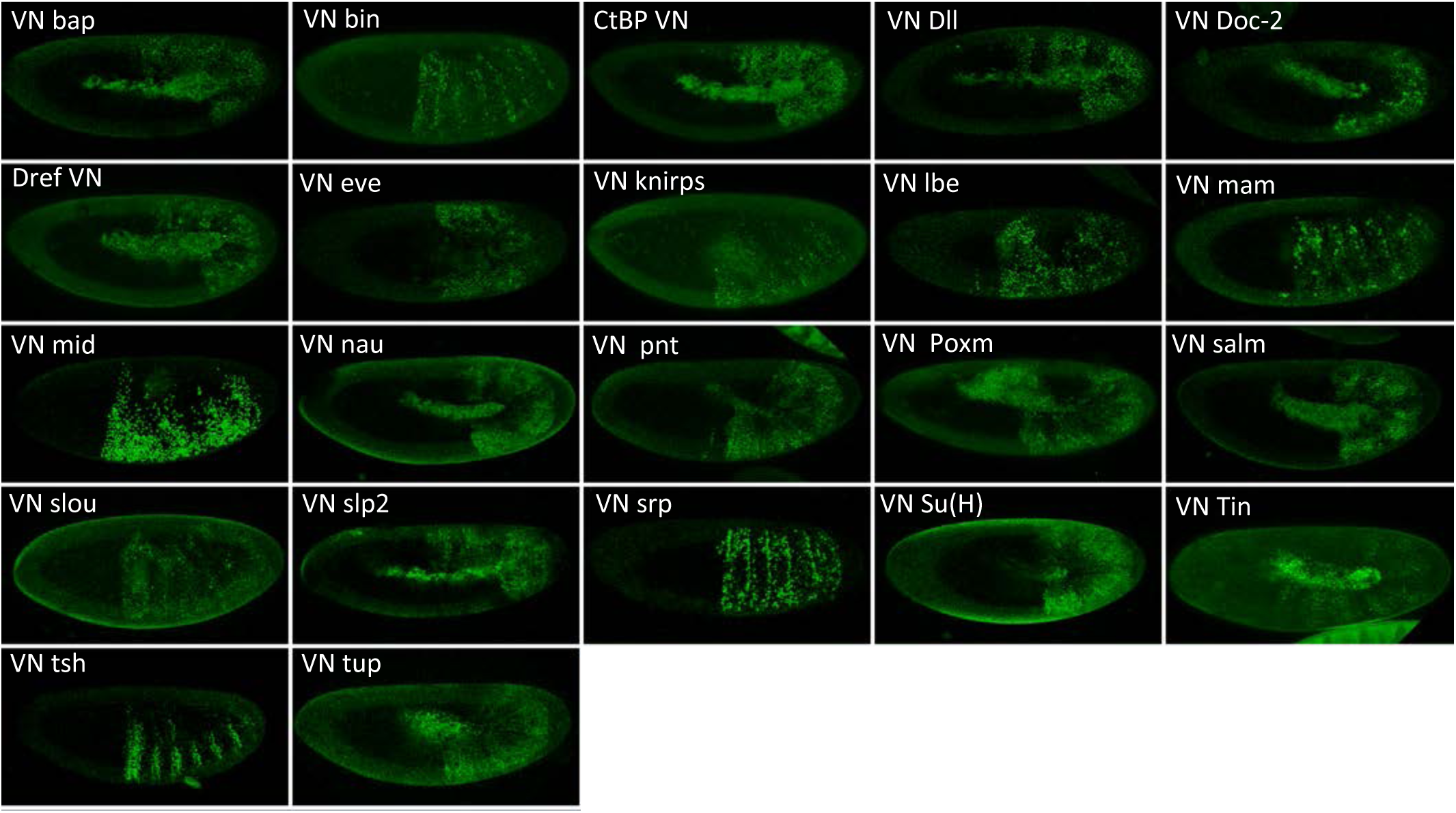
Illustrative confocal pictures of BiFC between ORF-VN and VC-Exd, as indicated. Fusion proteins are expressed with the *abdA-Gal4* driver and BiFC analysed in the epidermis of stage 10 embryos.

**Supplementary Figure 14.**
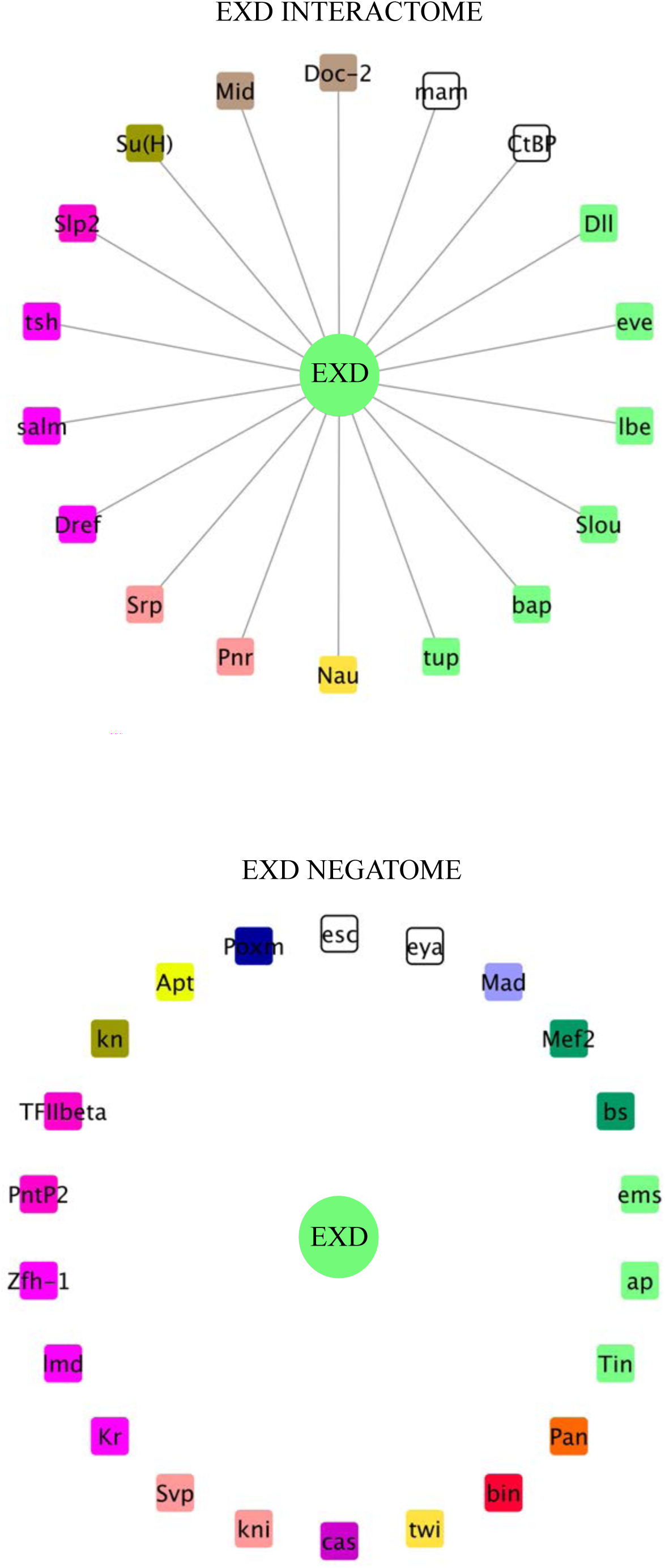
Interaction properties of Extradenticle (Exd) with a set of 37 TFs that are positive with AbdA. **A.** Representation of the interactome (positive interactions) of Exd. **B.** Representation of the negatome (negative interactions) of Exd.

## Supplementary Table legends

**Table S1. List of the 235 TFs of the multicolor BiFC library fused to the VN fragment at the C- or N-terminus, with a long or short linker region, as indicated.** Note that several TFs are fused to VN with either a long or a short linker region.

**Table S2. List of the 326 TFs of the multicolor BiFC library fused to the CC fragment.** TFs highlighted in light green are also available as VN fusion constructs (see Table S1).

**Tables S3. List of the 260 VN- and CC-fusion TFs of the multicolor BiFC library that were tested with Ubx and AbdA.**

**Table S4. Expression profile of the 260 TFs that were tested with Ubx and AbdA among 25 different developmental contexts of the *Drosophila* embryo.** Each color code corresponds to a different tissue when expressed. Black boxes depict expression in tissues where Ubx and AbdA are not present. The last two columns indicate the number of cooccurrences of the TF and the Hox protein with regard to the total distribution (parentheses).

**Table S5. Tandem comparison of Ubx- and AbdA-specific interactomes with regard to their respective enrichment in SLiMs, ordered and disordered regions.** Green and red boxes respectively highlight a significant enrichment or depletion (the corresponding p values are given between parentheses).

**Table S6. Analysis of the proportion of AbdA- and Ubx-specific TFs expressed in the different tissues, as indicated.**

**Table S7. List of TFs tested in RNAi in Ubx heterozygous mutant haltere discs.** Green and red boxes correspond to increased or not increased RNAi phenotypes, respectively. Black boxes correspond to RNAi phenotypes that could not be interpreted with regard to a potential Ubx cofactor function in the haltere disc (due to morphogenesis defects). Stars indicate RNAi fly lines that were tested with dicer. Stocks without the star are from the last generation and were tested without dicer.

**Table S8. List of VN fusion TFs that were positive in BiFC tests with VC-AbdA and tested with VC-Exd.** Green and red boxes indicate a positive or negative interaction status, respectively.

**Table S9. List of the 35 TFs tested as VN or CC fusion constructs with VC-AbdA or VN-AbdA, respectively.** Yellow boxes highlight the two TFs that showed opposite interaction status in the context of the two different fusion topologies (Hr83 and Ravus).

## Acknowledgements

We thank the Bloomington stock center for fly lines and Bart Deplancke for sharing the TF ORF plasmid library and Cristina Bastos for assistance with injections. Work in the laboratory of S. Merabet was supported by Association pour la Recherche sur le Cancer (ARC, PJA20141202007), Fondation pour la Recherche Médicale (FRM, DEQ20170336732), Ligue Nationale Contre le Cancer, Centre National de Recherche Scientifique (CNRS), CEFIPRA (5503-2), and Ecole Normale Supérieure (ENS) de Lyon.

